# Time calibrated tree of *Dioscorea* (Dioscoreaceae) indicate four origins of yams in the Neotropics since the Eocene

**DOI:** 10.1101/224790

**Authors:** Ricardo S. Couto, Aline C. Martins, Mônica Bolson, Rosana C. Lopes, Eric C. Smidt, João M. A. Braga

## Abstract

The yam genus *Dioscorea* comprises circa 650 species of tropical vines with starch rich tubers, usefull as an energy source and often containing secondary metabolites. The Neotropical Region holds the highest diversity of species and morphology of yams. We generated a time-calibrated tree for *Dioscorea* using, for the first time, a dense sampling of Neotropical species (64 sp., 20% of all Neotropical sp. and 22 sections) to trace the biogeography of these plants in this region. Four origins of *Dioscorea* in the neotropics were estimated since the Eocene. The two most diverse lineages originated between the Eocene and Oligocene, respectively in the Southern Andes and eastern South America. Both lineages occupied the South American ‘Dry Diagonal’ after the Miocene, but New World II clade remained associated with forest habitats. Several exchanges between Dry Diagonal and adjacent forested biomes occurred, corroborating the interchange between these vegetation types. Dispersals to Central America occurred before the closure of the Panama Isthmus. We highlight two important events of long distance dispersal, the colonization of Central American before the closure of Isthmus of Panama and the dispersal of *D. antaly* lineage to Madagascar. In addition, our phylogenetic tree evidenced the unnatural nature of the classical infrageneric classification of *Dioscorea*. The taxonomic implications of our results are also discussed.

## INTRODUCTION

*Dioscorea* L. comprises approximately 95% of the known species of Dioscoreaceae (Govaerts, Wilkin, & Saunders, 2007), as a reflection of this larger number of species the genus possess a tremendous morphological diversity, as several critical traits to distinguish from the other genus of the family (eg.: hermaphroditism in the other genera and dioecism in *Dioscorea*), and a considerable richness of chemical and genetic characters. Most species of *Dioscorea* are known as yam (and variations in different languages: inhame, ñame, igname, niam, enyame, nyami, etc.). They are mostly dioecious vines, usually with small flowers and starch-rich tubers as the underground organ. Due to the great nutritional value of its underground system, they are an important source of energy for human populations worldwide (Wilkin *et al*., 2005). *Dioscorea* species have their history linked to humans for thousands of years by a slow and gradual process of domestication (Ayensu & Coursey, 1972). In addition, *Dioscorea* is economically important due to secondary metabolites present in the tubers; *diosgenin,* for example, has a molecular structure similar to steroidal hormones and has been used to synthesize steroids for the production of birth control pills, among other uses (Coursey, 1967). Yams have been studied by its secondary metabolites, however studies related on prospection of useful substances in tubers of Neotropical species are restricted to only a few species (eg.: *D. composita, D. floribunda* and *D. mexicana*) even with the great diversity of neotropical species (Mignouna, Abang, & Geeta, 2009; Coursey, 1967).

The family Dioscoreaceae currently includes 653 accepted species, distributed in four genera: *Dioscorea* (633 species), *Stenomeris* Planch. (2 species), *Tacca* J.R. Forst. & G. Forst. (17 species), and *Trichopus* Gaertn. (1 species) (Govaerts, Wilkin, & Saunders, 2007). A total of 1600 names are attributed to *Dioscorea,* among species, varieties and subspecies, mostly considered as synonyms (The Plant List, 2013). *Dioscorea* species occur mostly in tropical areas with some representatives in subtropical and temperate regions of the planet, but are especially diverse in the Neotropics, where about 50% of the species occur (e-Monocot team, 2017).

According to Viruel *et al*. (2016) in its study with 135 taxa and four plastid DNA markers, *Dioscorea* originated in the Laurasian Palearctic region between the late Cretaceous (57.7 – 85.9 mya) and the Mid Eocene (47.6 – 49.1 mya), with subsequent radiations to the Southern regions by long-distance dispersal or migration by land bridges in the Oligocene-Miocene (33.9 to 5.332 Mya) (Viruel *et al*., 2016). They occur in several Neotropical environments, from dry *restinga* at sea level to the Andean paramos, including edges and interior of humid forests, natural grassland ecosystems, rupicolous areas, and even semi-desertic environments (Dorr & Stergios, 2003; Couto *et al*., 2014). As a consequence of the great variety of environmental conditions in which they occur, *Dioscorea* species exhibit a wide range of ecological responses, evidenced by the large morphological variability found in the family, both in vegetative and reproductive organs. They range from large climbing vines (40m high) to dwarf species, monoecious or dioecious plants, and they can present impressively colored leaves and flowers, among other distinctive characters (Fig. 1).

**Fig. 1.**
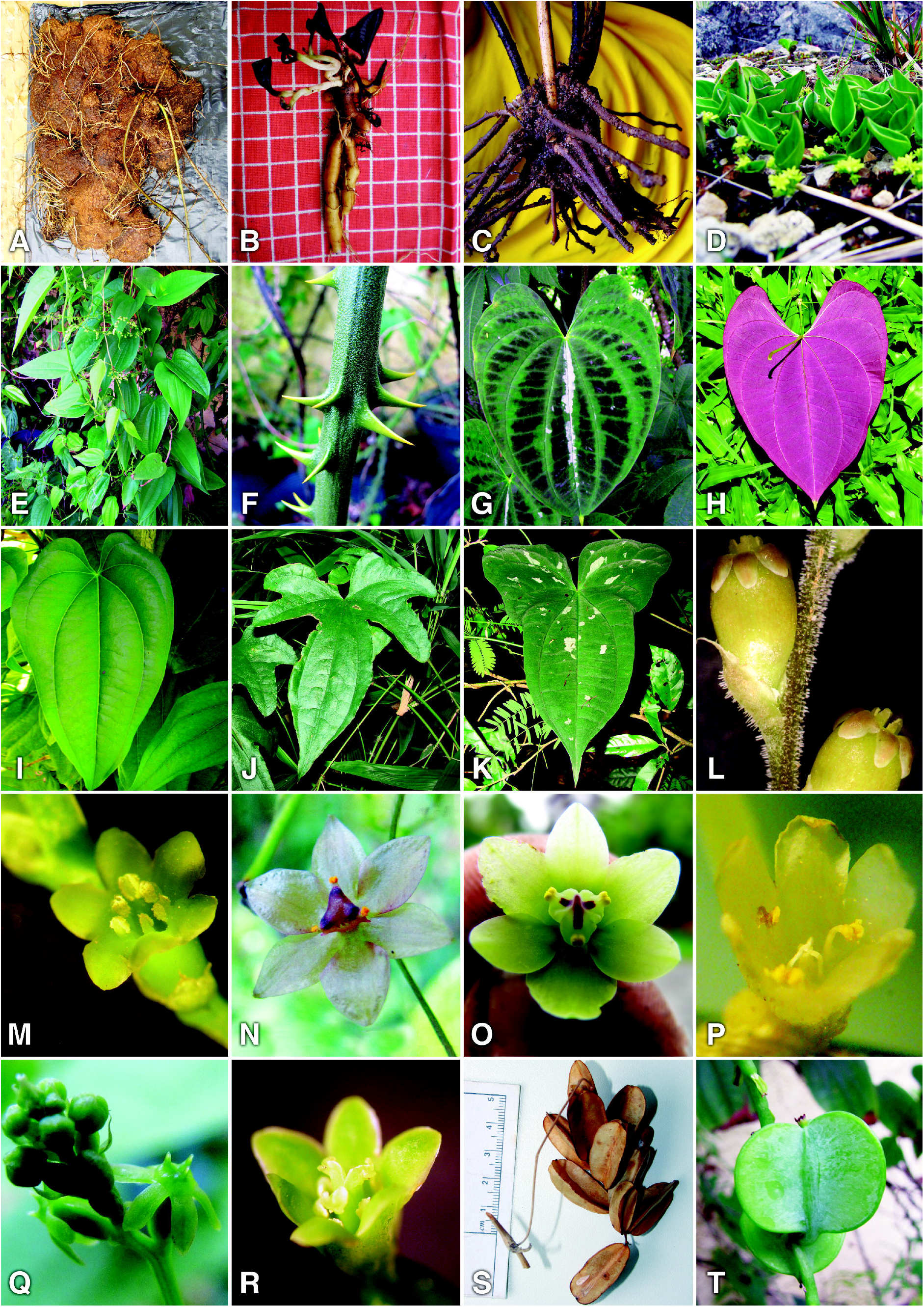
Neotropical *Dioscorea* morphological diversity. **a** tuber of *D. therezopolensis*. **b** tuber of *D. perdicum*. **c** underground organ of *D. multiflora*. **d** dwarf plant of *D. perdicum*. **e** habit of *D. campos-portoi*. **f** spines of *D. mollis*. **g-h** colorful leaves of *D. dodecaneura*. **i** leaves of *D. pseudomacrocapsa*. **j** lobeded leaves of *D. sinuata*. **k**. lobaded and variegated leaves of *D. hassleriana*. **l** staminate flower of *D. asperula*. **m** staminate flower of *D. laxiflora*. **n** staminate flower of *D. demourae*. **o** staminate flower of *D. subhastata*. **p** staminate flower of *D. sinuata*. **q** pistilate flower of *D. monadelpha*. **r** pistilate flower of *D. laxiflora*. **s** fruit of *D. subhastata*. **t** fruit of *D. olfersiana*.

Knuth (1924) proposed 58 sections and four subgenera for *Dioscorea: D*. subgenus *Helmia* (Kunth) Uline (seeds winged basally), *D*. subgenus *Dioscorea* (Pax) Uline (seeds winged all around), *D*. subgenus *Stenophora* (Uline) Knuth (seeds winged apically) and *D*. subgenus *Testudinaria* (Salisb.) Uline (seeds winged apically). This classification has undergone subsequent changes and additions (Burkill, 1960; Barroso *et al*., 1974), although these were not based on phylogenetic studies. With great morphological variation and scarce knowledge on anatomy, ecology, chemistry and palynology of *Dioscorea* (Ayensu, 1972; Caddick *et al*., 1998; Schols *et al*., 2001, 2003; Wilkin *et al*., 2009), external morphology was the base for the delimitation of taxonomic groups. Recent phylogenetic studies based on molecular data suggest eight main lineages distributed in the tropical and subtropical regions of the globe: ‘*Enantiophyllum*’, ‘Compound Leaved’, ‘Malagasy’, ‘Birmanica’, ‘Africa’, ‘European’, ‘New World’ and ‘*Stenophora*’ (Wilkin *et al*., 2005; Viruel *et al*., 2016). Viruel *et al*. (2016) obtained some clades not found in previous analyses: *Dioscorea* section *Rajania* (L.) Raz, a group of species endemic to the Caribbean islands (Raz, 2016), the clade *Epipetrum,* proposed by Philippi (1864) as genus for dwarf-sized species endemic to Chile (Viruel *et al*., 2010), and the clade *Nanarepenta,* for non-winged seeds species exclusive to Mexico, also previously proposed as a separate genus (Téllez-Valdés & Dávila-Aranda, 1998). Those clades fund by (Viruel *et al*., 2016) were sampled by previus studies (Caddick *et al*., 2002a,b; Wilkin *et al*., 2005)(), but (Viruel *et al*., 2016)presented a expanded the number of species and more a robustly supported results.

Full understanding of relationships among species of *Dioscorea* and a natural infrageneric classification will only be possible with strongly supported phylogenetic studies. However, the phylogenetic results to date do not allow a new formal and complete classification, especially for the Neotropical region, which is still poorly sampled in most recent published works (five neotropical taxa in Caddick *et al*., 2002a); three in Caddick *et al*., 2002b; 43 in Viruel *et al*., 2016). It should also be noted that the genera mentioned above — which are now known to be part of *Dioscorea* — still require a change of status regarding their positioning in an infrageneric taxonomic category within *Dioscorea*, with exception of *Rajania* that has been reduced to a section by Raz (2016).

Previous phylogenetic studies points to three events of radiation of *Dioscorea* into the Neotropics (Wilkin *et al*., 2005; Viruel *et al*., 2016), however their sampling did not completely reflect the diversity of Neotropical region. Species of *Dioscorea* arrived at least thrice in the Neotropics since the Oligocene and greatly diversified there, occupying various habitats in this region (Viruel *et al*., 2016). Since previous phylogenies did not include a dense sampling of Neotropical species, much of the taxonomic, morphological and geographical variation of Neotropical *Dioscorea* has not been covered, hampering inferences regarding its origin and diversification in this biogeographical region.

The present study increases by 45% the sampling of Neotropical species compared to previous phylogenetic studies in *Dioscorea*, including wider taxonomic and morphological diversity, and summed with sequences available in GenBank (mostly from Wilkin *et al*. (2005) and Viruel *et al*. (2016)), it represents the most densely sampled phylogeny of *Dioscorea* to date. This neotropical sampling (64 species in total) represents around 20% of all neotropical species. In terms of possible neotropical lineages, Knuth (1924) presented 56 sections for *D*. sect. *Helmia* and *D*. sect. *Dioscorea* (17 for *Helmia* and 39 for *Dioscorea*), 38 of these feature neotropical distribution (67%, 13 for *Helmia* and 23 for *Dioscorea*). In our study, we cover 22 of those sections (almost 40% of all sections, 7 for *Helmia* and 15 for *Dioscorea*) and four more species *Incertae sedis,* on which 14 of these lineages (10 sections and four *Incertae sedis*) had not been sampled in previous studies.

By reconstructing the phylogeny of *Dioscorea*, with special addition of Neotropical samples, we aimed to infer phylogenetic relationships among Neotropical taxa, as well as their arrival and divergence times in this region. As the richest lineages of the genus, we seek to test the infrageneric classification proposed Knuth (1924) (reestablishing any of the sections if supported by phylogeny) and to obtain a clear view of the lineages existing now at the Neotropical region, especially in cases where the Knuth, (1924) classification does not present clear delimitations between sections. Also, the understanding of how they occur in past, up to the present environments can provide tools for a better comprehension of the evolution of this group so rich morphologically. We also offer a more robust scenario of phylogenetic relationships in the genus, facilitating future proposals of an updated classification of *Dioscorea*.

## MATERIALS AND METHODS

### Taxon sampling, DNA sequencing and alignment

Species sampling aimed to cover a wide geographic and morphological range within *Dioscorea*. We produced new sequences for 34 species of *Dioscorea*: 12 spp. representatives of D. subg. *Helmia,* 18 spp. of *D*. subg. *Dioscorea,* and four spp. *incertae sedis,* plus *Tacca artocarpifolia* Seem as an outgroup representative of Dioscoreaceae (Table S1). The taxon sampling aimed to cover a great part of the lineages proposed as sections by classic authors (sensus the revisionmade by Knuth, 1924), covering 22 of the 38 sections proposed to the neotropical region, attending the sections with great number of species with more than one taxa (eg.: *D*. section *Dematostemon* Griseb., five species sampled in diferent morphological strata).

Most new sequences were generated from specimens collected in the field by R. S. Couto, prioritizing taxonomically well-delimited species from well-known populations. Vouchers were deposited in the Herbarium of National Museum (R), with duplicates at the Herbarium of the Botanical Garden of Rio de Janeiro, Brazil (RB).

Additionally, we included 143 sequences from GenBank: 129 from Dioscoreaceae (119 species of *Dioscorea,* two species *Trichopus,* seven of *Tacca* and one *Stenomeris*), and five from other families (three species of Burmanniaceae, one Stemonaceae and one Thismiaceae), totaliing 177 species sampled. All sampled species with geographic origin, herbarium vouchers and GenBank accession numbers are listed in Table S2.

Total genomic DNA was extracted from leaf samples, fresh or silica-dried following the 2x CTAB protocol (Doyle & Doyle, 1987), without the addition of RNase A and scaled to 2 ml tubes. The extracted DNA was measured in 1% agarose gel with a DNA mass ladder and deposited at the DNA collection of Laboratory of Systematics and Molecular Ecology of Plants, Federal University of Paraná (UFPR), Brazil, associated to the reference vouchers deposited in K, R, RB, RFA and UPCB (Herbarium acronyms follow *Index Herbariorum* (Thiers, continuously updated)). The plastid genome regions mat*K* and rbc*L* were amplified and sequenced using the following primer pairs, respectively: 3F_KIM-f (cgtacagtacttttgtgtttacgag) and 1R_KIM-r (acccagtccatctggaaatcttggttc) (Ki-Joong Kim, pers. com.), rbcLa_f (atgtcaccacaaacagagactaaagc) (Levin *et al*., 2003) and rbcLa_r (gtaaaatcaagtccaccaccrcg) (Kress & Erickson, 2007). Additionally, the primers ITS92 and ITS75 or ITS18F and ITS26R, which is widely used in angiosperms (Bolson *et al*., 2015), was used to amplify the nuclear ITS region, however without sucesss in sequencing. This negative result and the scarcity of ITS sequences for *Dioscorea* in GenBank lead us to use only plastidial genome markers. PCR amplifications were performed using initial 94°C pre-melt for 1 min followed by 40 cycles of (i) 94°C denaturation for 30s, (ii) 53°C annealing for 40s, and (iii) 72°C extension for 40s, followed by 72°C a final extension for 5 min. Following PCR, the samples were purified with 20% PEG and sequenced with Big Dye Terminator version 3.1 (Applied Biosystems, California, USA) by the company Macrogen Inc. (South Korea). Forward and reverse sequences were assembled using the Staden package v.2.0.0b11 (Staden, Judge, & Bonfield, 2003). Sequences were aligned with Clustal W using default parameters (Thompson *et al*., 1997) implemented in the software MEGA6 (Tamura *et al*., 2013).

### Geographical and morphological data

We examined over 4,000 specimens deposited in 79 herbaria, in addition to field observations, with special effort on the Neotropics, for the selection of taxons sequenced, to obtain morphological comparisons between the analyzed species (detailed in the discussion of the clades) and especially for coding the geographic distributions used in biogeography. Specimens from the following herbaria were examined: B, BAA, BAFC, BR, C, CAY, CEPEC, CESJ, COAH, COL, CR, CTES, CUVC, CVRD, ESA, F, FAA, FCAB, FURB, GUA, HAL, HAS, HB, HCF, HMUC, HRCB, HST, HUCP, HUEFS, HUPG, HVASF, HXBH, IAC, ICN, INPA, IPA, IRBR, JE, JVR, K, L, LPS, M, MBM, MEXU, MG, MNHN, MO, MVFA, MVFQ, MVM, NY, OPUR, P, PACA, PEL, PH, R, RB, RBR, RFA, RFFP, S, SI, SMDB, SP, SSUC, U, UFP, UFPR, ULS, UNR, UPCB, US, UV, WU, XAL, Z, and ZT. Herbarium acronyms follow Index Herbariorum (Thiers, continuously updated).

### Phylogenetic analysis

Maximum Likelihood, Parsimony and Bayesian inference were used to estimate tree topologies on the concatenated matrix of 177 taxa and 1658 nucleotides from rbc*L* and mat*K* genes. Maximum-likelihood tree searches were performed using raxmlGUI v.1.0 (Silvestro & Michalak, 2012) under the model GTR+I+G and statistical support for nodes were assessed with 1,000 bootstrap replicates, consistent with that used by Viruel *et al*. (2016). Parsimony analyses were conducted in PAUP 4.0b10a (Swofford, 2002) using heuristic tree searches with tree bisection-reconnection (TBR), 2,000 random-taxon-addition replicates holding 20 trees per replicate. Branch support was estimated with 2,000 bootstrap pseudo-replicates (Felsenstein, 1985). Bayesian phylogenetic inference with the Metropolis-coupled Markov Chain Monte Carlo (MCMC) was used to estimate tree topology and posterior probability distribution as implemented in MrBayes v.3.3.4 (Ronquist *et al*., 2012), using two parallel runs, each with four chains and 10 million generations, with parameters sampled every 1,000 generations. Each gene was considered as one partition and the best-fitting models under the Akaike Information Criterion in the software Mega 6 (Tamura *et al*., 2013) were GTR+G (mat*K*) and K2+G (rbc*L*). Convergence of runs was assessed in Tracer 1.6 (Rambaut, Suchard, & Drummond, 2014). A 25% burnin was applied to eliminate trees prior to convergence of chains and a 50% majority rule consensus tree was constructed from the remaining trees.

### Molecular dating

The *Dioscorea* tree molecular clock was estimated using fossil and secondary calibrations and the full matrix of 177 taxa including 162 representatives of *Dioscorea* and 15 outgroups. The fossil record of Dioscoreaceae was recently reviewed by Raz (2017), who analyzed twenty fossils attributed to this family from different time periods and geographical origins, mostly leaves. Only three can be attributed to sections within *Dioscorea* and therefore are more suitable for molecular dating: *Dioscoroides lyelli* (Eocene), *Dioscorea wilkinii* (Oligocene) and *Dioscorea* sp. from Kenya (Sect. Asterotricha) (Miocene). Other fossils are attributed only to genus level or would require further information to confirm their position, and eight of them are not Dioscoreaceae (Raz, 2017). Three Dioscoreaceae fossils used to calibrate the tree were chosen by stratigraphic reliability and confidence of taxonomic assignment (Raz, 2017). We follow the fossil calibration described by Viruel et al. (2016): A. The fossilized leaf of *Dioscoroides lyelli* from the Eocene at the Paris basin, France (Potonié, 1921), dated from the Ypresian age, was assigned to the *Stenophora* stem due to similarities with the extant *Stenophora* species; B. The fossil seed attributed to *Tacca buzekii* from the Upper Eocene from Putschirn, Czech Republic (Gregor, 1983), was assigned to the crown node of *Tacca*; C. The fossilized leaflet attributed to *Dioscorea wilkinii,* from the Middle Oligoceneof Ethiopia (Pan, Jacobs, & Currano, 2014), was assigned to the crown node of section *D*. Section *Lasiophytum* A lognormal distribution prior was applied to all fossil calibrations nodes, with values also following Viruel *et al*. (2016), as follows: *Dioscoroides lyelli* (mean= 48.2, sd= 0.008); *Tacca* (mean: 35.85, sd= 0.028); *Dioscorea wilkinni* (mean: 27.23, sd= 0.002). Calibration points are depicted in Fig. S3.

Divergence times were inferred under a relaxed uncorrelated lognormal clock model in BEAST 1.8.3 (Drummond *et al*., 2012) implemented in the CIPRES server (Miller, Pfeiffer, & Schwartz, 2010), using a Yule tree model of speciation, and HKY+I+G substitution model with empirical base frequencies. The MCMC chains ran for 50 million generations, sampled every 10,000 generations. Convergence was assessed using effective sample size (ESS) values ≥ 200 in Tracer 1.6 (Rambaut *et al*., 2014). Two separated runs were performed and their results were combined in LogCombiner, totalizing 100 million generations. The maximum clade credibility tree was generated in TreeAnnotator (BEAST package), and visualized and edited in FigTree v 1.4 (Rambaut, 2009).

### Biogeographic analysis

In order to estimate ancestral distribution areas of Neotropical *Dioscorea* we defined six major geographic areas: A= Central America, B= Northern Andes, C= Southern Andes, D= Amazonia, E= Dry Diagonal and F= Atlantic Forest. Definitions were based on the current distribution of Neotropical species of *Dioscorea*, considering areas with more than 10 endemic species, seeking to exclude areas that present only occasional endemisms. The delimited areas were also based on the Neotropical regional classification proposed by Morrone (2014) and a study on Rubiaceae by Antonelli *et. al*. (2009). Although another six species that also occur in the Neotropics were included in our analysis, we focused our biogeographical analysis on the most diverse clades, New World I and New World II (hereafter NWI and NWII).

We estimated ancestral range probabilities on the multimodel approach performed by the R package BioGeoBEARS (Matzke, 2013; R Core Team, 2016) using the three available models: DEC, BAYAREALIKE and DIVALIKE. The dispersal-extinction-cladogenesis model (DEC) considered cladogenetic processes as the evolution of range at speciation events and allows the estimation of free parameters *d* (dispersal) or range extension and *e* (extinction) or range loss by maximum likelihood (Ree & Smith, 2008). Dispersal-vicariance-analysis (DIVA) is a parsimony-based method that allows dispersal and extinction in anagenetic processes and vicariance in cladogenetic processes (Ronquist, 1997). The model is called DIVALIKE in BioGeoBEARS because it is a maximum likelihood implementation of DIVA. The BayArea method is a Bayesian approach specifically designed to analyze a large number of areas efficiently (in reasonable computer time) (Landis *et al*., 2013). BAYAREALIKE implemented in BioGeoBEARS is a maximum likelihood interpretation of BayArea. The founder speciation event parameter *j* was also added to all analyses, creating the models DEC+*j*, DIVALIKE+*j*, BAYAREALIKE+*j* The parameter *j* adds the possibility of a new cladogenesis event, where an individual ‘jumps’ to an area completely outside the ancestral range, founding a new genetically isolated lineage (Matzke, 2013, 2014). For selection of best-fit model, we relied on the best likelihood value as well as the Akaike Information Criterion (ΔAICc).

## RESULTS

### Phylogenetic analysis

*Dioscorea* species form a strongly supported clade (bootstrap support value (BS) ML and MP 100%, Posterior Probability (PP) 1) (Fig. 2, Fig. S1 and S2). Among the internal clades, *Dioscorea* section *Stenophora* has strong support (BS-ML and BS-MP 100%, PP 1) and represents a sister lineage to the remaining *Dioscorea*. Traditional classification systems and phylogenetic analysis results are compared in Table S4.

**Fig. 2.**
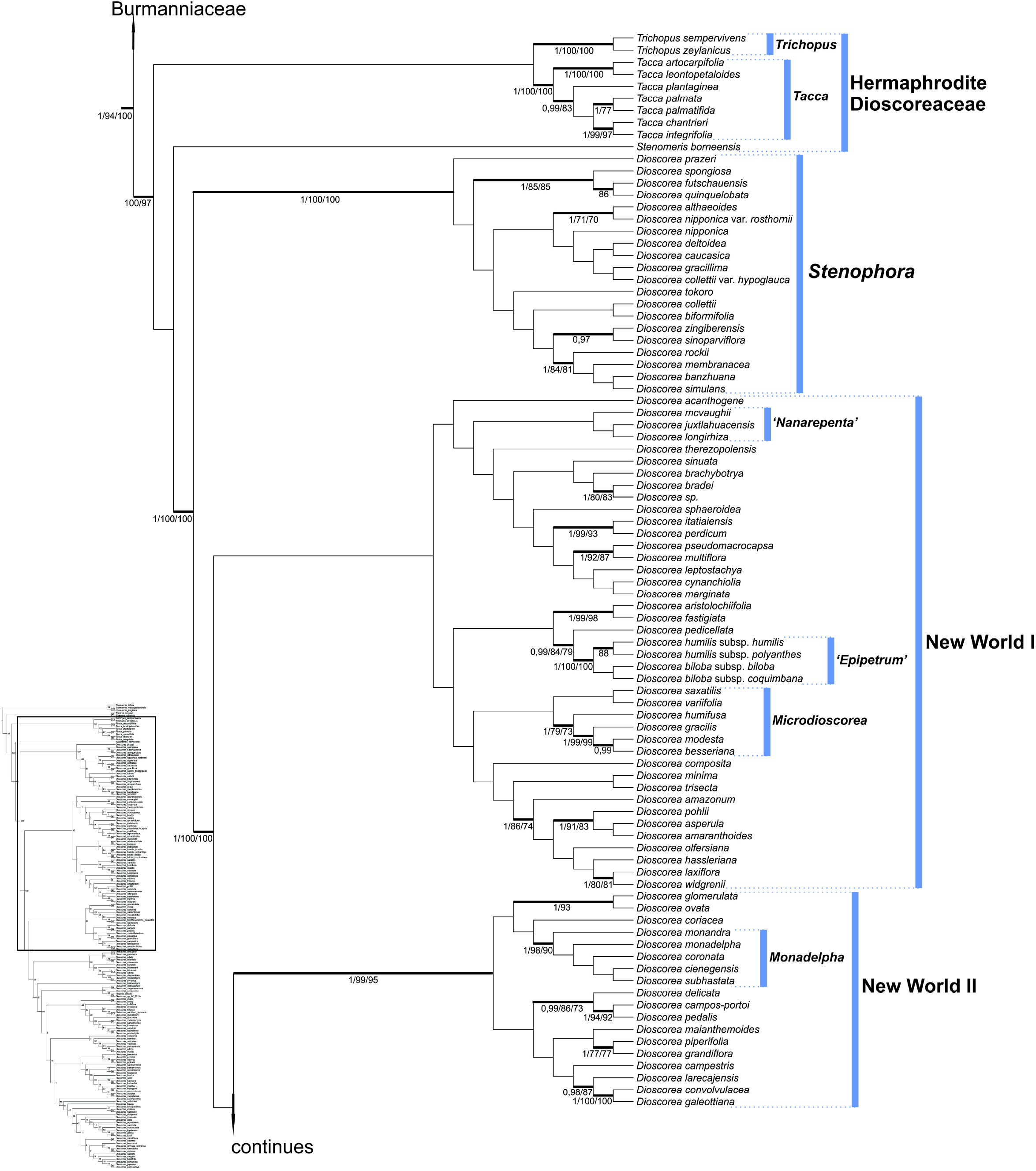
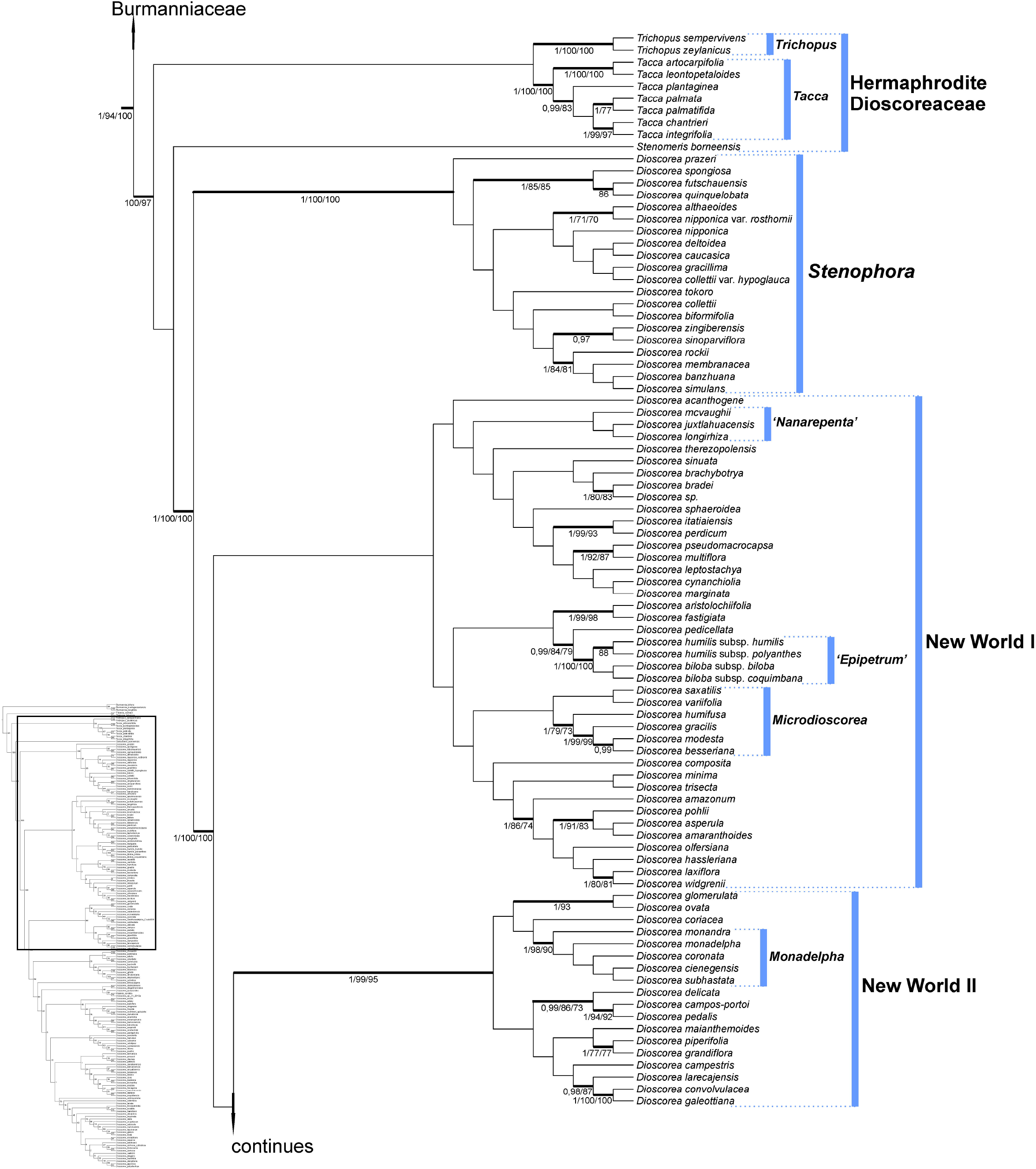
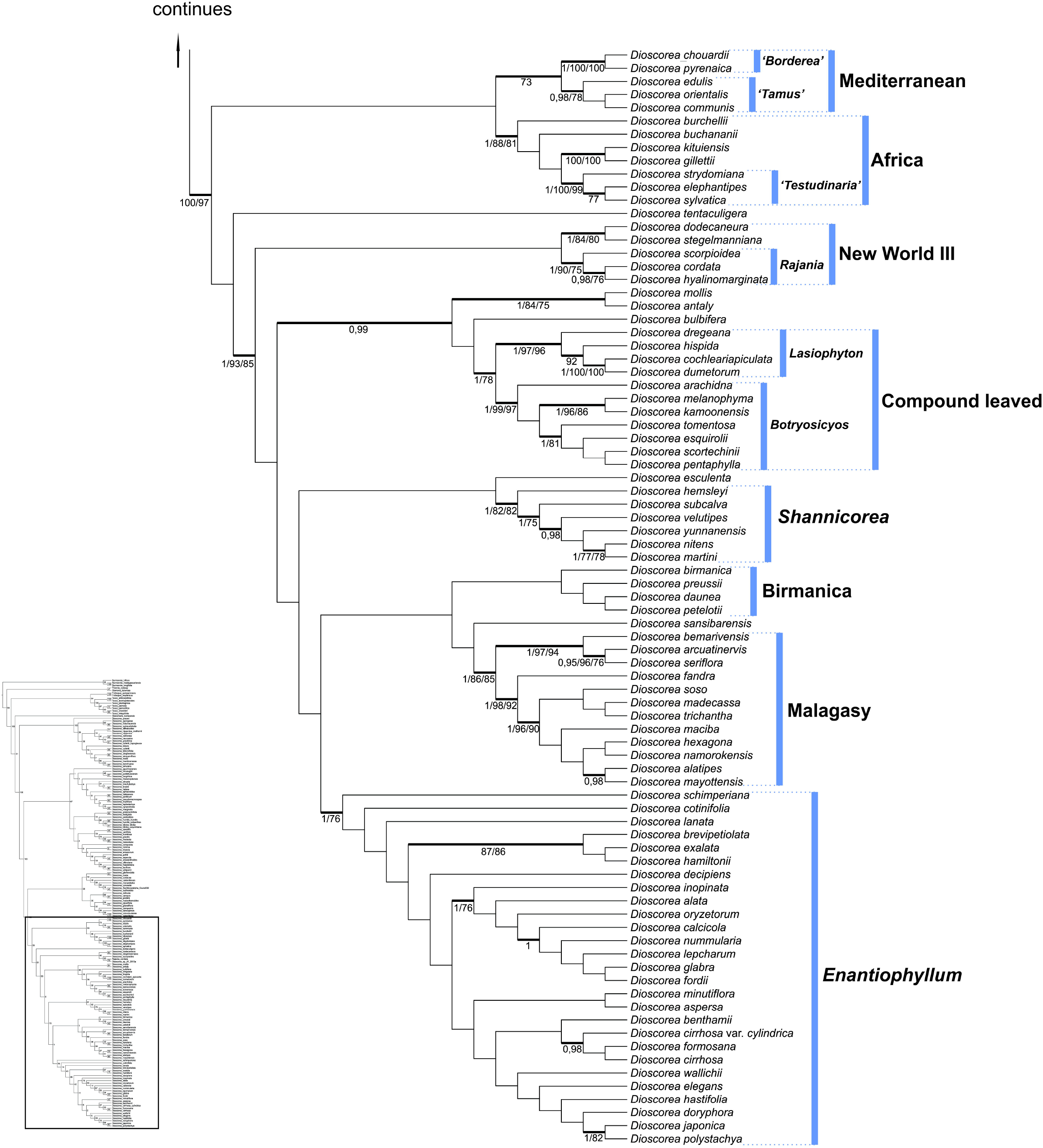
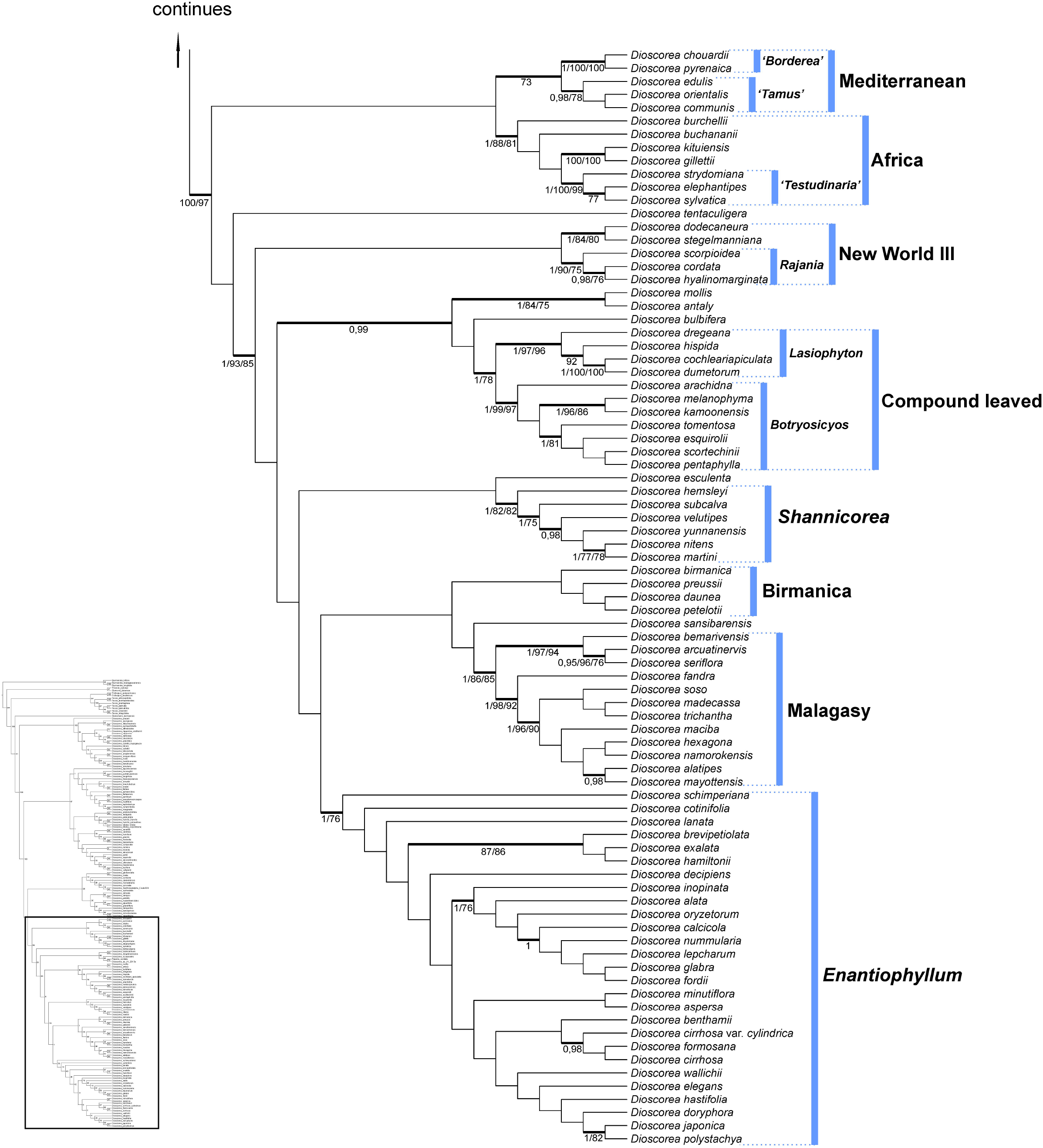
Maximum likelihood tree derived from the analysis of 177 taxa and 1658 nucleotides of *Dioscorea* and outgroups. Thickened lines represent highly supported branches in at least one of the three analysis (ML, BI, and MP). Values on nodes represent supports obtained in the three analyses, respectively: BI, ML, MP. Branches with bootstrap values □70% and BPPs □95□ were considered weakly supported.

Most Neotropical species grouped into two clades, here indicated as ‘New World I’ and ‘New World II’. New World I groups the species corresponding to *D*. subg. *Dioscorea,* restricted to the Neotropics. Within this clade it is possible to recognize another three small clades: *Epipetrum,* with high support (BS ML and MP 100%, PP 1), *Microdioscorea,* and *Nanarepenta,* with low support. New World II has strong support (BS-ML 99%, BS-MP 95%, PP 1) and groups species corresponding to *D*. subg. *Helmia* restricted to the Neotropics. The *Monadelpha* clade can be recognized in the New World II clade, as a strongly supported clade (BS-ML 98%, BS-MP 90%, PP 1) with sister species. Neotropical species that were external to NWI and NWII, *D. dodecaneura* Vell., *D. stegelmanniana* R.Knuth, and species representatives of *D*. section *Rajania* (L.) Raz form the clade ‘New World III’, appearing associated with clades of African and Asian species (Africa and Compound Leaved). The species of *D*. section *Rajania,* however, form a well supported clade (BS-ML 90%, BS-MP 75%, PP 1) within New World III. Only one species with Neotropical occurrence appears in this analysis outside the clades mentioned above, *D. mollis,* which is sister to *D. antaly,* and related to Asian species.

The clade *Shannicorea* appears for the first time and brings together the seven species from Southeast Asia, with moderate support (BS ML and MP 81%, PP 1). The remaining Old World species are organized in a large clade, where the inner clades ‘Mediterranean’, ‘Africa’, ‘Compound Leaved’, ‘Birmanica’, ‘Malagasy’ and *Enantiophyllum* can be highlighted, following Viruel *et al*. (2016) nomenclature. The results obtained in this analysis are congruent with Viruel *et al*. (2016), just with some differences in the support values of the clades, those values can be retrieved in Figure 1 and others in supplementary material (Fig. S1, S2, S3).

Bayesian and Maximum Likelihood analysis results did not differ considerably, except for small differences in the most recent clades in ‘Birmanica’, *D*. section *Shannicorea* and ‘Malagasy’. Maximum Parsimony analysis also resulted in high support for most of the clades that were well-supported in other analyses (Fig S2).

### Divergence times and biogeography

According to our divergence time analysis in BEAST, the most recent common ancestor of Dioscoreaceae (including *Tacca*) originated in the Cretaceous around 99 Mya (95% Highest posterior density interval (HPD): 72.7–132.4 Mya) (Fig. 3). All divergences in the genus level occurred in the Paleocene. The most diverse genus in the family, *Dioscorea*, is estimated to have originated in the Cretaceous-Paleocene boundary, around 66 Mya (54.8 –84.6 Mya) (stem age). The two main Neotropical clades originated between the Eocene and Oligocene: the crown age for New World I is 37.2 Mya (28–44.3 Mya) and around 28 Mya (20.8–37.8 Mya) for New World II. The third Neotropical clade comprises only four species and originated in the Oligocene, around 30 Mya (20.3 –37.5 Mya). The clade that includes Neotropical *Dioscorea mollis* plus the Malagasy *D. antaly* originated in the Miocene at 18.5 Mya (6–31.5 Mya).

**Fig. 3.**
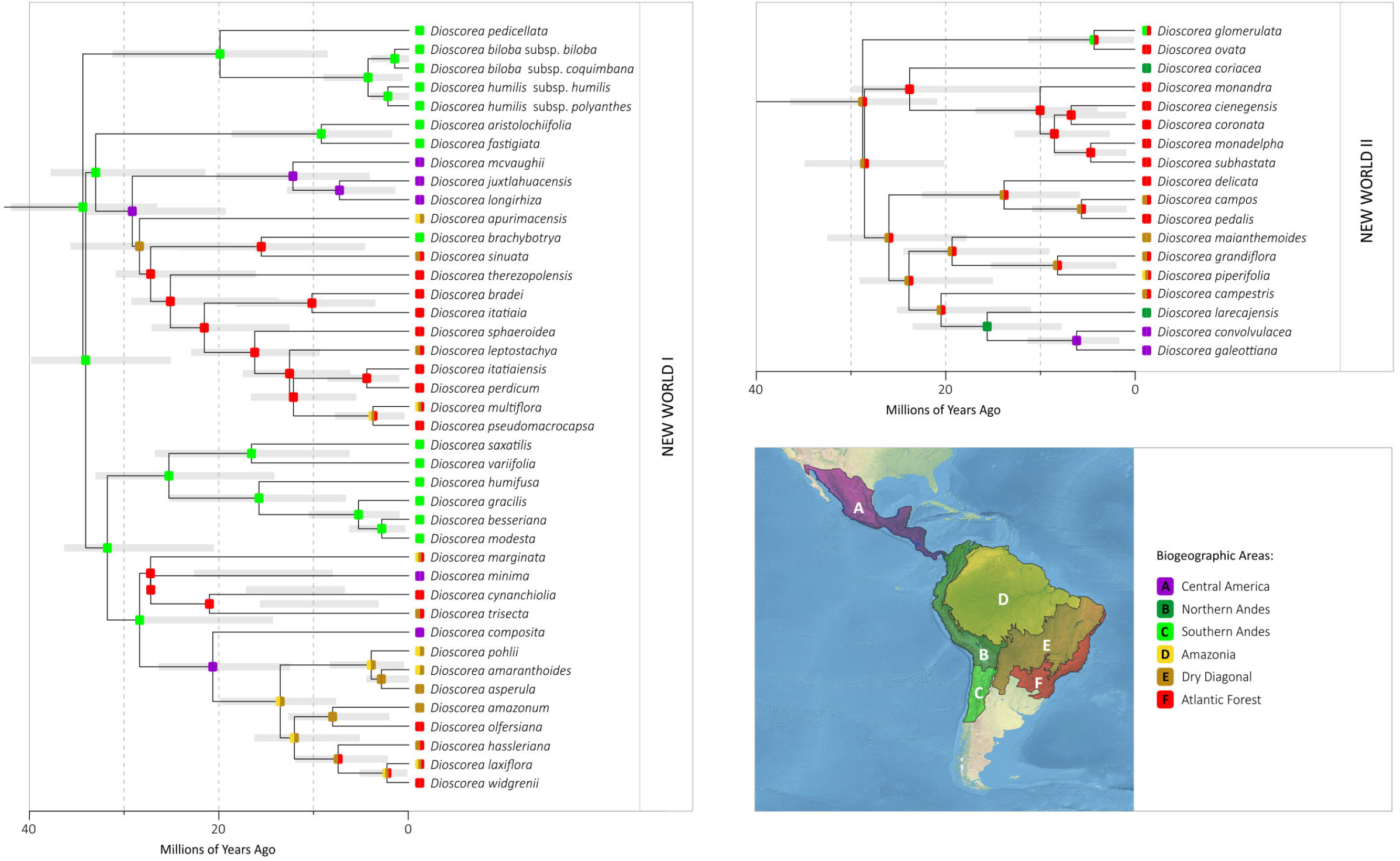
Combined time-calibrated tree and ancestral area reconstruction analyses for the groups New World I and II of *Dioscorea*. The tree is the maximum clade credibility tree based on the BEAST analysis of the molecular matrix with 177 taxa and 1658 nucleotides for *Dioscorea* and outgroups calibrated using fossils. Grey bars on nodes indicate the 95% confidence interval. Diagrams on nodes show the single most-probable ancestral range using the best model BAYAREALIKE+j in BioGeoBEARS (See Table S3 for the summary statistics). Squares on the tips represents the present range of each terminal. Areas used in the biogeographic analysis: A= Central America, B= Northern Andes, C= Southern Andes, D= Amazonia, E= Dry Diagonal and F= Atlantic Forest.

Biogeographic analyses performed separately on clades New World I and New World II yielded the same model as the best scored model (i.e. with lowest likelihood): BAYAREALIKE+j (See Table S3 for all parameter results). The ancestral distribution of the MRCA for New World I clade in *Dioscorea* is predicted to be the “Southern Andes” biogeographic region (Fig. 3), a region that nowadays comprises Northern Argentina and Chile and corresponds to the southern part of the mountain range, not present at the time of origin of NWI clade. Some clades remain endemic to this area (for example, the *D. fastigiata* – *D. humilis* clade). *Dioscorea* expanded its distribution eastward during the Oligocene, diversifying in today’s Atlantic Forest. Expansions to Central America probably occurred in the Miocene at least twice. Further occupations of Dry Diagonal plus Amazonia and/or Atlantic Forest occurred in the Middle Miocene (around 10 Mya).

In contrast, New World II is estimated to have originated in the eastern part of South America, i.e. in a region comprising the Dry Diagonal and Atlantic Forest (Fig. 3). Most of the extant species in this clade still inhabit these regions. In the Miocene, two independent occupations of Northern Andes occurred, corresponding to the *D. coriacea* and *D. larecajensis* stems. Approximately at the same time, around 15 Mya, the *D. convolvulacea* – *D. galeottiana* lineage dispersed into Central America, with *D. convolvulacea* still being distribuited in Northern South America.

Two species of very restricted distribution in completely distinct regions appear in a small recent clade in the phylogenetic analysis of *Dioscorea*, *D. mollis* and *D*. *antaly*, which are endemic species of Southeastern South America (Atlantic Forest) and Madagascar (Tropophylles Forest = Deciduous Dry Forest), respectively. *Dioscorea antaly* is the only endemic species of Madagascar to appear outside the Malagasy clade, and *D. mollis* is the only endemic species of the Neotropics to emerge outside the New World clades (NWI, NWII and NWIII).

## DISCUSSION

### Phylogenetic analysis and classification of *Dioscorea* and position of Neotropical clades

Eleven major clades were obtained for the monophyletic genus *Dioscorea*, which was consistent with the clades observed in other recent studies. The nine clades and sections of *Dioscorea* already established in other studies (Wilkin *et al*., 2005; Viruel *et al*., 2016) were also recovered in this analysis, with the emergence of internal clades such as *D*. section *Shannicorea* (even though weakly supported) and the position of Neotropical species external to the clades New World I and II (Fig. 2).

The taxa of unique occurrence in the Neotropics form basically two clades, as also stated by Wilkin *et al*. (2005) and Viruel *et al*. (2016). The New World I clade (NWI) groups (although, with low support) the species with circular or semicircular winged seeds that occur in the tropical region of the Americas, once placed by Knuth (1924) in *D*. subgenus *Dioscorea* along with other *Dioscorea* of circular or semicircular winged seeds with wider, pantropical distribution. The seed morphology is part of the *Dioscorea* taxonomy since the the first classifications proposed, Kunth (1850) proposed the *Helmia* genus using (beside others) this character, as other follow in theirs infrageneric classification. Although Knuth (1924) knew that the use of this character to split the *Dioscorea* genus almost at half was not completely adequate, nor reflected a systematic relationship, it was used, at least for practical reasons, for many years as the base of the infrageneric classification of *Dioscorea*. Wilkin *et al*. (2005) had already shown that this character and subgenus classification were not a reflection of the evolutionary relations of the group, however, they obtained several clades where one of the states of this character were fixed in all specis sampled in their analisys. The autors used this condition to help to explain and separate the especies in the two internal clades of Madagascar and as a plesiomorfic condition of all species of their clade “Compund-leafed”, yet they didn’t discussed this character in the scope of the “New World” clade (probably by the lack of knowledge of the fruit morphology of neotropical species at the time).

Allthough, this clade does not include all the species placed by Knuth (1924) in this subgenus of pantropical distribution, it contemplates species from different regions and habitats of the Neotropics, as well as great morphological diversity (Fig. 1). This could mean that this character, had only a few changes between the the two states, at least in the Neotropics, where almost half of all species of the genus are distributed, and could point to a probable sinapomorphy to the NW I and II clades.

The Chilean species in the NWI clade are basically arranged in three smaller clades (*Epipetrum, Microdioscorea* and *Nanarepenta,* with exception of *D. brachybotrya*), with poor internal resolution. In spite of the evidence for the relationship between the taxa endemic to Chile, the *Epipetrum* clade groups dwarf species that present non-winged seeds, characteristic to the dry regions of Chile, as seen in Viruel *et al*. (2016). Besides these species, we have six taxa placed by Knuth (1924) in the *D*. section *Microdioscorea,* a section composed mostly of Chilean species (only two taxa for Peru and Brazil) with stems of reduced length and six stamens. *Dioscorea* section *Microdioscorea* appears as monophyletic with low support in this analysis and also in Viruel *et al*. (2016).

Another clade within the NWI group is the group endemic to Mexico, previously designated by Matuda (1961) as the genus *Nanarepenta*, which also has non-winged seeds. In Viruel *et al*. (2016), *D. minima* appeared along with the other three species of *Nanarepenta,* but in the present analysis this species is distantly related. This lack of resolution could be a reflection of the increased number of taxon and the lower amount of markers used in this study. Due to the great diversity found in the Neotropics is desirable that a greater number of species, as well as the DNA regions, could be analysed to solve the internal relations of the largest Neotropical clade.

Whithin the species of NWI clade is possible to highlight *D. perdicum* Taub., one of the dwarf species endemic from Brazil that does not fit any specific infrageneric classification devotaded to this issue. *Dioscorea perdicum* is mistakenly placed by Knuth (1924) into the section *Cycladenium* Uline (*D*. subg. *Helmia*), as the autor didn’t know the species fruits or seeds, that place *D. perdicum* in *D*. subg. *Dioscorea* by its all-round winged seeds. Burkill (1960) pointed out the mistake made in the allocation in the *Cycladenium* section suggesting to be placed in *D*. sect. *Pedicellatae* R. Knuth, but most importante he proposed this species as a possible link between the Neotropical and Paleotropical species, with relation to *D*. sect. *Stenophora*, specifically by the presence of what he called a rhizome. As a matter of fact, the underground organ of *D. perdicum* is a tuber of rhizomatous aspect (Fig. 2), as have all anatomical features of a tuber (Tenorio, pers. comm.) but the aspect is similar to a rhizome, as doesn’t have cataphylls but it branches and produces roots and shoots from more than one point. This unique morphology leaded Burkill (1960) to assume the close relation to the Paleotropical species of *Dioscorea*, aspect not confirmed in our analisys, as *D. perdicum* appears without any special relation to the *Stenophora* clade.

The other species that are endemic to the Neotropics appear here as the strongly supported clade NWII (BS-ML 99%, BS-MP 95%, PP 1), which contains part of the formerly designated *D*. subg. *Helmia* (sensu Knuth, 1924), unlike the NWI clade, this group has been retrieved in all phylogenetic studys that have sampled the neotropical species (Wilkin *et al*., 2005; Viruel *et al*., 2016) and also strog support, demonstrating the likely single origin of the elongated seed wings in the Neotropical region. Even though the sections sampled here have been increased only by one, the number of species within key sections (*D*. sect. *Dematostemon* Griseb. and *Monadelpha* Uline) has been increased in order to contemplate more comprehensively the diversity of this group. *Dioscorea* sect. *Dematostemon* is one of the biggest sections of the Neotropical region, comprising 45 taxons of great morphological diversity and high distribution range, going from dwarf species (eg.: *D. anomala* (Kunth) Griseb. and *D. maianthemoides* Uline ex R. Knuth) endemic to the brazilian “Cerrado” to typical Atlantic Forest species *(D. campanulata* Uline ex R. Knuth and *D. cinnamomifolia* Hook.). Knuth (1924) also placed in this section species (D. *moyobambensis* R. Knuth, *D. galiiflora* R. Knuth and *D. triangularis* (Griseb.) R. Knuth) closely related to others in *D*. sect. *Centrostemon* Griseb., showing that the boundaries of this group are not well defined. With our increased sampled analisys, this section is shown to be polyphyletic.

In addition, we have a small clade with strong support, formed by *D*. section *Monadelpha* species, which present unusual characteristics, such large staminate flowers, three fertile stamens forming a fleshy column, free and entire stylus in pistillate flowers. Besides those characteristics, this section possess an almost unique feature related to sexuality in the genus, the monoecy, all species from this section present staminated inflorescences in the lower part of the plant and pistillated on the higher part. This character is only shared with a few species from *D*. section *Cycladenium* (probably misplaced) and *D. margarethia* G.M. Barroso, E.F. Guim. & Sucre (poorly know compoud-leafed species with six stamens on a column instead of three), and besides some unstable dioecy in *D*. sect. *Stenophora* and *D. convovulacea* Schltdl. & Cham. (Wilkin *et al*., 2005). The monoecious species from *D*. section *Cycladenium* aro not well know and the boundaries of the section are not clear, as is a section with great diversity (i.e. monoecious and dioecious species). In our analisys, the only species from *D*. section *Cycladenium, D. coriacea* Humb. & Bonpl. ex Wild., (dioecious), doesn’t appear to be closely related to this clade. Addition of more taxa from this group could clarify relationships the two sections and reinforce the monophyly of *D*. section *Monadelpha*.

The *Monadelpha* clade also appear as monophyletic in other studies with less dense taxon sampling for the Neotropic (Wilkin *et al*., 2005; Viruel *et al*., 2016), however, with no mention of the section, due in part to the smaller sample of the *Monadelpha* clade and due to a misidentification of one of the species used in Viruel *et al*. (2016) (D. *monandra* Hauman was iderntified as *D. calderillensis,* from *D*. section *Cycladenium*). Therefore, we understand that monoecy probably originated only once in the in the family, with origin in the Neotropical region. The position of *D. margarethia* should be tested to reinforce this organization, but the species is only know from its type specimens and a second gathering (that could not be sequenced), and even the fact that it has six stamens doesn’t seem to be a problem as this character is volatile in the NW clades.

Even with the increased number of taxa sampled to the NWI clade from 23 to 64 species and 15 Knuth’s sections for *D*. subg. *Dioscorea* and seven for *D*. subg. *Helmia,* plus four *incertae sedis* species the resolution in the Neotropical clades is not yet close to be solved. It has been increased the number of internal clades monophyletic (*Monadelpha, Nanarepenta, Epipetrum, Microdioscorea*) and species of most sections of Knuth (1924) are shown not to be phylogenetically related (e.g.: *Dematostemon, Apodostemon* Uline, *Cryptantha* Uline).

A small group of Neotropical species emerged outside the main Neotropical clades (NWI and NWII), being composed of three species from *D*. section *Rajania,* plus two South-American species, *D. dodecaneura* and *D. stegelmanniana*. Wilkin *et al*. (2005) already presented *Dioscorea cordata* (L.) Raz as a distinct lineage, separate from the NWI and NWII clades, which can also be observed in Viruel *et al*. (2016), also associated to the neotropical edible species *D. trifida* L. f.. Those species have very different morphological characteristics when compared to other Neotropical species, some of these characteristics being shared with those found in species from Asia and Africa. The common ground to this species is the presence of annualy renewed tubers, those are the only neotropical species sampled so far that have this character (absent in the NW clades and rather rare in the neotropical species). The matter of annual tubers has been addressed by Wilkin *et al*. (2005), showing that is a paleotropical characteristic (only present in their B clade), being these clades the ones related here to NWIII.

*Dioscorea dodecaneura* and *D. stegelmanniana* are morphologically very similar to each other, but they present marked differences in one key aspect of Knuth’s classification, the fruit (transversely oblong and oblong, respectively) and seeds shape (circular and oblong, respectively). This indicates that besides the NWI and NWII clades the seed wing shape is not stable character, having closely related species on the NWIII clade with both of the states of this characteristic. Additionally, these two species present a particular pattern of organization of the vascular bundles of the aerial stem (Tenorio *et al*., 2017), similar to that described by Ayensu (1972) as the typical Old World pattern. It is noteworthy that *D. trifida* and *D. stagelmanniana* have been sorted by Knuth (1924) to *D*. sect. *Macrogynodium* Uline, reflecting in his view the close relationship of this species.

*Dioscorea* section *Rajania* is composed of 18 species, besides one non-described species, with occurrence restricted to the West Indies (Raz, 2016; Raz & Pérez-Camacho, 2016)(Raz, 2016). The species belonging to *D*. section *Rajania* are distinguished by the samaroid fruits, although this is not an exclusive feature of this section, as pointed out by Raz (2016) in the most recent taxonomic treatment on this group. The morphological characteristics exhibited by this section exemplify the diversity found in the Neotropical region, even though it is a clade with lower morphological diversity, it presents great differences for the rest of the neotropical species and with a more recent arrival when compared to the other two Neotropical clades (NWI and NWII), with the maintenance of characteristics typical of species of the Asian and African region.

*Dioscorea mollis* shares several characteristics with paleotropical species, such as phyllotaxis ranging from alternate to subopposite or even opposite, a characteristic found in less than 2% of the species of the American continent, where most species presents alternate leaves. The species also present an underground system composed of several fibrous nodules from which numerous aerial stems appear (similar to a rhizomatous system), stems of woody aspect and prickles in the basal stem, which are also unusual characteristics for Neotropical species. These characteristics are shared with one closely related species of *D*. section *Chondrocarpa* Uline, *D. chondrocarpa* Griseb., not sampled here by the unsuccessful amplification of *matK* gene. *Dioscorea chondrocarpa* was also sampled by Viruel et al. (2016) but did not reached the final publication by lack of genes amplified successfully, however its position in their inicial analisys topology is congruent to the one fund here to *D. mollis* (Raz, pers comm.). Anatomically these species also presents similarities to the Old World species, as verified by Tenorio *et al*. (2017). All these evidences strongly indicate a fourth lineage of *Dioscorea* in the Neotropical region, more related to paleotropical species.

The Paleotropical clades obtained in our analysis were similar as those recovered in previous studies (Wilkin *et al*., 2005; Maurin *et al*., 2016; Viruel *et al*., 2016), consisting in the Africa clade with the inner clade *Testudinaria* (composed of species occurring in the mountainous regions of eastern and southern Africa), the Malagasy clade (with all endemic species from the island of Madagascar, except for *D. antaly*), and the clade *Enantiophyllum* (composed by several species proposed to the *D*. section *Enantiophyllum* Uline). Previous and the present phylogenetic results contradict the main infrageneric classifications of *Dioscorea* (Uline, 1897; Knuth, 1924; Burkill, 1960), which grouped the species in various sections. Some aspects of those clade of these clades are interesting to emphasize, such as the Malagasy clade is internally organized into two small clades, one presenting circular winged seeds, and the second grouping the remainder species with elongated winged seeds, and that *Enantiophyllum* (with its enormous diversity and imprecise delimitation) has some polytomies in the present and previous phylogenetic analyses, indicating the need of more data to elucidate the internal relationships

*Dioscorea* section *Shannicorea* was proposed to group six species of occurrence restricted to Asia, mostly China. The taxa share the left twining stem, the staminate inflorescences composed of small scorpioid cymes, stamens inserted at the base of the tube segments and the elongated seed wings. Knuth (1924) treats the same species, with the addition of two taxa, as *D*. section *Shannicorea,* but subordinated to *D*. subg. *Stenophora*. In contrast to the initial position proposed by Uline (1897) for the *Stenophora* section, Knuth (1924) elevated *Stenophora* to subgenus and further organized it internally into two sections, *Eustenophora* R.Knuth and *Shannicorea* Prain & Burkill.

In our analyses, the six species listed by Prain & Burkill (1914) in *D*. section *Shannicorea* (*D. hemsleyi, D. martini, D. nitens, D. subcalva, D. velutipes* and *D. yunnanensis*) are grouped in a single clade, presenting similar internal relationships in all analyses, but positioned differently in the *Dioscorea* tree. It is the first time that this clade is supported in a widely sampled *Dioscorea* tree. Hsu *et al*. (2013) also recovered this clade, but their sampling included only species from East and Southeast Asia. Viruel *et al*. (2016) also analysed some of these species (*D. nitens* and *D. subcalva*), retrieving them inside the Birmanica clade, as they are related. In our analysis, *D*. section *Shannicorea* is distantly related to *D*. section *Stenophora,* demonstrating that Knuth’s (1924) proposal to treat *D*. section *Shannicorea* as part of the subgenus *Stenophora* has no phylogenetic support and Prain & Burkill’s (1914) proposal could be more accurate. This indicates that the section as proposed by Prain & Burkill (1914) could be monophyletic, but more DNA markers in a phylogeny that includes species from both clade that is closely related (*Shannicorea* and Birmanica) are needed to solve uncertainties in this part of the topology.

The placement of *D. sansibarensis* at the base of the Malagasy clade raises some questions regarding the evolution of this group in an insular environment. According to Viruel *et al*. (2016), Madagascar was colonized by *Dioscorea* species from Asia, and not from Africa as many angiosperms. On the other hand, the sister species *D. sansibarensis* occurs in several areas of Africa, besides Madagascar, presenting a high vegetative dispersal ability: they massively produce small aerial tubers in the leaf axils, which possibly facilitated their invasive behavior in several countries (Raz, 2002; Choo, 2009; Hsu & Wang, 2012). The presence of *D. sansibarensis* in Madagascar could be product of a recent natural dispersal event or human introduction, since this species has been used for food and for the production of venom (Wilkin *et al*., 2005).

We have new evidence suggesting that within both the biggest New World clades, some of the sections proposed by Uline (1897) may be supported and can fit in a phylogenetic sound revised classification of *Dioscorea* to come. The position and monophyletism of *D*. section *Microdioscorea* and *D*. section *Monadelpha* within a widely-sampled phylogenetic analysis of *Dioscorea* shows that the proposed infrageneric classification by classical works (Kunth 1924; Uline 1897) mostly do not represent natural lineages, but some of them still may be used in modern systematics of the genus. Increasing the Neotropical species sampling evidenced the role of this group of species as key to provide a complete and accurate infrageneric classification of *Dioscorea,* as it is the most diverse and taxonomically complex region, and at the same time, the most underrepresented in phylogenetic studies until the present study. Nevertheless, the Neotropical species still lack a wider sampling to reach a better resolution of these clades, 16 sections of the Knuth (1924) classification still don’t have been used in any phylogeny up to the present (22 of 38 were coverd here), and this should be goal to persue to better undertand the infrageneric classification of Neotropical *Dioscorea*.

### *Dioscorea* lineages originated four times independently in the Neotropics since the Eocene

The phylogenetic analysis presented here, focused on the Neotropical clades of *Dioscorea,* provides a new perspective on the biogeographical history of this genus in South and Central America. The biogeographic analysis has shown that four independent lineages of *Dioscorea* diversified into the Neotropical Region, two of them becoming highly diverse and wide spread. The Neotropical species of *Dioscorea* present at least four different origins. The New World I and II clades are more diversified and widely distributed (Fig. 2), while another two species are grouped in the predominantly Caribbean group *Rajania,* and *D. mollis* is sister to the Malagasy *D. antaly*. With the exception of *D. mollis,* which has an independent origin, all the other hypotheses on origins of Neotropical clades of *Dioscorea* had been described before (Viruel *et al*., 2016). For the first time, however, we presented a more detailed view on the biogeographic history of the group after the colonization of the Neotropics.

The pantropical genus *Dioscorea* putatively originated in Laurasia during the Late Cretaceous – Early Eocene, later dispersing into South America, Africa and Madagascar (Viruel *et al*., 2016). The colonization of America could have been facilitated by the existence of land bridges during the Palaeocene-Eocene thermal maximum (Zachos, Dickens, & Zeebe, 2008), such as the North Atlantic Land Bridge (NALB) and Beringean Land Bridge (BLB), presumably during early Oligocene (Viruel *et al*., 2016), further reaching South America by occasional island chains such as the proto-Greater Antilles (Antonelli *et al*., 2009). Nevertheless, ancestral area reconstruction suggests a South American origin for the Neotropical *Dioscorea* with further dispersals towards Central America (Fig. 3), a result that was also found in the global analysis by Viruel *et al*., (2016). Exchanges between Laurasia and South America are reported for other plant groups, such as Malpighiaceae, which is similarly pantropical, greatly diverse in South America, and possibly migrated through Laurasia after having originated in South America (Davis *et al*., 2002).

At least three different origins are estimated for the Neotropical clades. The NWI clade originated in the Eocene-Oligocene boundary, in what is now the Southern Andes Region, and dispersed to eastern South America and Central America. At this time most of the Andes were not formed yet, which could allow eastwards expansions of range; however, the South American continent was partially occupied by marine incursions from the Caribbean and from the Pacific seas, which could be barriers to expansions (Antonelli *et al*., 2009). Expansions towards the East in South America occurred only after the Oligocene, around 30 Mya. Since the end of Cretaceous, the South American humid forests dominated the terrestrial habitats and no evidence of dry vegetation exists for this period until the decrease in temperatures that took place after the Miocene Medium Climatic Optimum (Davis *et al*., 2005; Hoorn *et al*., 2010).

Occupation of dry vegetation biomes occurred in both clades, but only sparsely in NW II, which remained almost exclusive to rainforests. The MRCA of the clade that occurs in Dry Diagonal is ambiguous and could have been Amazonian or from the Dry Diagonal. It is known that the origin of the South American Dry Diagonal must have taken place only after 10 Mya, with climate gradual cooling and drying in the Miocene, and before the establishment of rainforests (Simon *et al*., 2009). Some species of *Dioscorea* occur in both open vegetation formations (Dry Diagonal) and forested biomes, i.e. Amazonia (*D. acanthogene, D. pohlii* and *D. amaranthoides*) and Atlantic Forest (D. *sinuata, D. leptostachya* and *D. trisecta*). The most generalist species can occur in the Dry Diagonal and the two forested biomes (*D. multiflora*, *D. marginata* and *D. laxiflora*). Species with occurrence in forested habitats usually will appear in arboreous vegetation patches in the Cerrado, like the “Cerradão” and Gallery forests or Caatinga forest, and not in open vegetation. Connections between Amazonia and Cerrado occurred many times in history, not only because of the geographic proximity of the regions occupied by the two vegetation types, but also because during episodes of climatic fluctuations, forests are known to have expanded or retracted (Costa, 2003). Quaternary cooling and drying episodes during the glacial times favored the expansions of savanna-type vegetation (Cerrado), decreasing the extent of tropical forests (Werneck *et al*., 2012). Only NWII presents species with distribution restricted to the open field formations (*D. maianthemoides*) and rocky savannas (*D. campos-portoi*) within the Cerrado biome.

Dispersals towards Central America occurred in Middle Miocene in both NW I and II, much earlier the estimated closure of the Ishtmus of Panama (3 Mya), which linked the still “isolated” biota of South America to North America (for a review of the long process of formation of Isthmus of Panama see O’Dea *et al*., 2016). Flora exchange before the Isthmus formation is best explained by long distance dispersals (Antonelli & Sanmartín, 2011; Freitas *et al*., 2016) and this seems to be the case of *Dioscorea*. Transoceanic dispersals explained much of plant biogeographic patterns and is hypothetically more explanatory than plate tectonics (Renner, 2004; Christenhusz & Chase, 2013). Besides the plant’s dispersal capability, which in the case of *Dioscorea* is facilitated by seed and fruit morphology and production of aerial tubers, wind and sea currents also facilitate the dispersal across oceans, generating a “dispersal pattern” (Renner, 2004). Many plant and animal groups dispersed from South to North America (or the opposite direction) starting in the Eocene, such as Malpighiaceae (Davis *et al*., 2002), *Hedyosmum* (Antonelli & Sanmartín, 2011) and Rubiaceae (Antonelli *et al*., 2009).

The sister group relationship between the Neotropical *D. mollis* and Malagasy *D. antaly* is quite unusual, however there are a few examples of sister clades occurring in Neotropical region and Africa or Madagascar in ferns, such as *Leucotrichum* (Polypodiaceae) (for more examples see Rouhan *et al*., 2012). Fern spores are efficient dispersal agents and greatly facilitated trans-atlantic dispersals and colonization (Rouhan *et al*., 2012). In Solanaceae, the genus *Tsoala* also dispersed from South America to Madagascar, probably by long distance dispersal facilitated by sea currents (Olmstead, 2013). In this family, most fruits are fleshy, animal dispersed, but many are dry and could have been dispersed by wind and sea currents, and dispersal occurred in both cases, but more frequently in the fleshy fruited lineages (Olmstead, 2013; Dupin *et al*., 2017). *Dioscorea antaly* could have dispersed from South America to Madagascar (or in the opposite direction) facilitated by its anemocoric dispersion syndrome, and specially by the shape of the seed, that is winged towards the base of the capsule, being more effective in high speed winds (Maurin *et al*., 2016). In gramitid ferns and *Tsoala* the long distance dispersal event occurred from the Neotropics towards Madagascar. Stem anatomy results indicated the proximity between *D. mollis* and other Neotropical species (Tenorio *et al*., 2017), possibly indicating that the direction of dispersal could have been from the Neotropical Region to Madagascar. Further phylogenetic analysis including more Neotropical species morphologically similar to *D. mollis* could further test this relationship hypothesis and clarify a scenario of dispersal between Neotropical Region and Madagascar.

## ACKNOWLEDGEMENTS

We would like to thank the Coordenação de Aperfeiçoamento de Pessoal de Nível Superior (Capes) for the research grants of RSC and ACM. ECS thanks Conselho Nacional de Desenvolvimento Científico e Tecnológico (CNPq) for Bolsa de Produtividade em Pesquisa CNPq-Nível 2 (processo 311001/2014-9). We would like to acknowledge Sylvain Razafimandimbison, for the contribuitions concerning his knowledge of Madagascar and its biogeography, Anamaria Dal Molin, for the revision of the English, Gabriel Rezende, for the preparation and revision of the figures, and the anonymous reviewers for they contribution on greatly improving the discussion of this paper.

## Supporting information

### Figures

**Fig. S1**. Bayesian consensus tree resulting from the analysis of the complete data set (177 taxa and 1658 nucleotides), rooted in Burmmaniaceae. The main clades of *Dioscorea* are highlighted. Posterior probability values >95 are shown on nodes.

**Fig. S2**. Maximum Parsimony tree resulting from the analysis of the complete data set (177 taxa and 1658 nucleotides), rooted in Burmmaniaceae. The main clades of *Dioscorea* are highlighted. Bootstrap values >70 are show on nodes.

**Fig. S3**. Bayesian maximum clade credibility time tree for *Dioscorea* and outgroups obtained under a relaxed clock model in BEAST and fossil calibration points. For all significantly supported nodes, bars show the 95% Highest Posterior Density intervals around the estimated ages. Fossil calibration points are: A. *Dioscorea lyelli* (Potonié, 1921); B. *Tacca* seed (Gregor, 1983); C. *Dioscorea wilkinii* (Pan *et al*., 2014).

### Tables

**Table S1.**
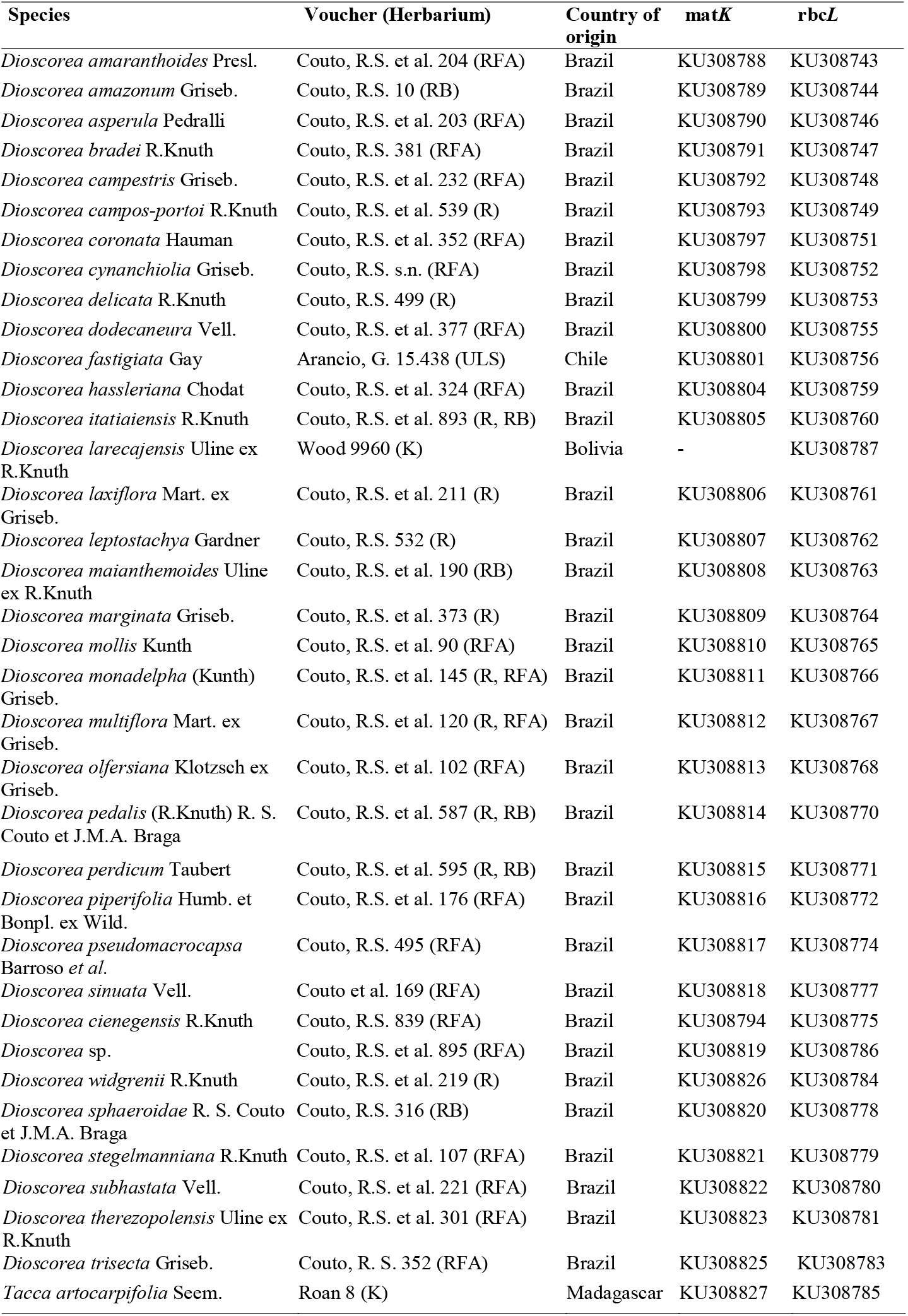
Sequences newly produced for this study, with information on voucher, country of origin and GenBank Accession Numbers. Herbarium acronyms follows the *Index Hebariorum* (Thiers, continuously updated).

**Table S2.** Species for which sequences were downloaded from GenBank with voucher information, GenBank accession numbers and references for the original publication. (Wilkin *et al*., 2005; Gao *et al*., 2008; China Plant BOL, 2011; Hsu *et al*., 2013; Viruel *et al*., 2016) (for complete reference, see References). Herbarium acronyms follows the *Index Hebariorum* (Thiers, continuously updated).

| Species | Voucher (Herbarium) | Reference | matk | rbcl |
| --- | --- | --- | --- | --- |
| *Burmanna biflora* L. | Chase 157 (NCU) | 1 | AY956483 | AF206742 |
| *Burmanna longifolia* Becc. | Johns et al. 9157 (K) | 1 | AY956484 | AF307484 |
| *Burmanna madagascariensis* Mart. & Zucc. | Caddick et al. 312 (K) | 1 | AY956485 | AF307486 |
| *Dioscorea acanthogene* Rusby | Lewis 3210 (LOJA) | 1 | AY956477 | AY667101 |
| *Dioscorea alata* L. | Wilkin et al. 1090 (K) | 5 | AB040208 | AY667098 |
| *Dioscorea alatipes* Burkill & H. Perrier | Caddick et al. 334 (K) | 1 | AY950687 | AY667099 |
| *Dioscorea althaeoides* R.Knuth | Zhu & Zhao 0648572 (NAS) | 2 | EU407548 | EU407550 |
| *Dioscorea antaly* Jum. & H.Perrier | Wilkin et al. 1104 (K) | 1 | AY956476 | AY667100 |
| *Dioscorea arachidna* Prain & Burkill | Wilkin et al. 862 (K) | 1 | AY956478 | AF307468 |
| *Dioscorea arcuatinervis* Hochr. | Caddick et al. 310 (K) | 1 | AY956479 | AY667102 |
| *Dioscorea aristolochiifolia* Poepp. | Villar & Viruel 20 (JACA) | 5 | KM877834 | KM877874 |
| *Dioscorea aspersa* Prain & Burkill | No data | 2 | EF614211 | EF614213 |
| *Dioscorea banzhuana* S.J. Pei & C.T.Ting | Huang & Guo 0648582 (NAS) | 2 | DQ974182 | DQ408174 |
| *Dioscorea bemarivensis* Jum. & H. Perrier | Wilkin et al. 1124 (K) e Caddick 340 (K) | 1 | AY956480 | AY667103 |
| *Dioscorea benthamii* Prain & Burkill | Venus 101 (TCB) | 4 | JQ733666 | JQ733743 |
| *Dioscorea bessleriana* Kunth | Villar & Viruel 14 (JACA) | 5 | KM877835 | KM877875 |
| *Dioscorea biformifolia* S.J. Pei & C. T. Ting | Zhu & Zhao 0648576 (NAS) | 2 | EU407549 | EU301740 |
| *Dioscorea biloba* (Phil.) Caddick & Wilkin subsp. *biloba* | Viruel & Villar Eb09 (JACA) | 5 | KM877836 | KM877876 |
| *Dioscorea biloba* subsp. *coquimbana* Viruel, Segarra-Moragues & Villar | Segarra-Moragues & Pérez-Collazos Eb01 (JACA) | 5 | KM877837 | KM877877 |
| *Dioscorea birmanica* Prain & Burkill | Thapayai et al. 409 (BKF) | 1 | AY956481 | AY667104 |
| *Dioscorea brachybotrya* Poepp. | Rudall 1/97 (K) | 1 | AY956482 | AF307469 |
| *Dioscorea brevipetiolata* Prain & Burkill | Wilkin et al. 964 (K) | 1 | AY956486 | AY667105 |
| *Dioscorea burchaninii* Benth. | Bingham et al. 10290 (K) | 1 | AY956487 | AY904790 |
| *Dioscorea bulbifera* L. | Foster et al. 12 (K) e NASHYY065 (200911007 KUN) | 1 | AY956488 | AY904791 |
| *Dioscorea burchellii* Baker | Wilkin s.n. SANBI DNA Bank MWC 33240 | 5 | KM877853 | KM877905 |
| *Dioscorea calcicola* Prain & Burkill | Wilkin et al. 814 (K) | 4 | AY956489 | JQ733799.1 |
| *Dioscorea monandra* Hauman | Wood et al. 22679 (K) | 5 | KM877849 | KM877891 |
| *Dioscorea caucasica* Lipsky | Zhou & Wu 0648584 (NAS) | 2 | DQ974188 | DQ408182 |
| *Dioscorea chouardii* Gaussen | Segarra-Moragues & Catalán s.n. (Silica specimen) | 5 | KM877855 | KM877907 |
| *Dioscorea cirrhosa* Lour. | Chen 51 (TCB) and Thapayai 375 (BKF) | 1 | JQ733663 | AY904792 |
| *Dioscorea cirrhosa* var. *cylindrica* C. T. Ting & M. C. Chang | No data | 5 | HQ637689 | DQ408184 |
| *Dioscorea cochleariopiculata* De Wild. | Chase 21046 (K) | 5 | KM877862 | KM877921 |
| *Dioscorea colletii* Hook.f. | Hsu 76 (TCB) | 4 | JQ733728 | JQ733805 |
| *Dioscorea colletii* var. *hypoglaucia* (Palib.) S. J. Pei & C. T. Ting | No data | 3 | HQ637622 | HQ637771 |
| *Dioscorea communis* (L.) Caddick & Wilkin | Sánchez de la Campa & Sánchez Civila s.n. (Silica specimen) | 5 | KM877854 | KM877906 |
| *Dioscorea composita* Hemsl. | No data | 2 | DQ974180 | DQ408172 |
| *Dioscorea convolvulacea* Schltldl. & Cham. | Chase 197 (NCU) | 1 | AY956491 | AJ235805 |
| *Dioscorea cordata* L. | Axelrod 8407 (UPR) | 1 | AY973835 | AF307472 |
| *Dioscorea coriacea* Humb. & Bonpl. ex Willd. | Lewis 3220 (LOJA) | 5 | AY956492 | KM877893 |
| *Dioscorea cotinifolia* Kunt | No data | 2 | DQ974185 | EF614219 |
| *Dioscorea daunea* Prain & Burkill | Thapayai et al. 518 (BKF) | 1 | AY956493 | AY904793 |
| *Dioscorea decipiens* Hook.f. | Wilkin et al. 861 (K) e Wilkin 860 (K) | 1 | AY956494 | AF307454 |
| *Dioscorea deltoidea* Wall. ex Griseb. | Huang & Guo 0648575 (NAS) | 2 | EF614207 | EF614218 |
| *Dioscorea doryphora* Hance | Hsu 56 (TCB) | 4 | JQ733669 | JQ733746 |
| *Dioscorea dregeana* (Kunth) T. Durand & Schinz | No data | 5 | KM877863 | KM877922 |
| *Dioscorea dumetorum* (Kunth) Pax | Wilkin 761 (K) | 1 | AY956495 | AF307464 |
| *Dioscorea edulis* Lowe | Catalán s.n. (Silica specimen) | 5 | KM877856 | KM877908 |
| *Dioscorea elegans* Ridl. ex Prain & Burkill | Rudall 113 (K) | 5 | KM877867 | KM877931 |
| *Dioscorea elephantipes* (L'Hér.) Engl. | UCI Arb. 6773 (NCU) | 1 | AY956496 | AF307461 |
| *Dioscorea esculenta* (Lour.) Burkill | Wilkin et al. 995 (K) | 1 | AY956497 | AY904794 |
| *Dioscorea esquirolii* Prain & Burkill | No data | 2 | DQ974177 | DQ408168 |
| *Dioscorea exalata* C.T. Ting & M.C. Chang | Y.S. Liang 1037 (TCB) | 4 | JQ733681 | JQ733758 |
| *Dioscorea fandra* H. Perrier | Caddick et al. 324 (K) | 1 | AY956498 | AY904795 |
| *Dioscorea fordii* Prain & Burkill | Shui 3526 (TNM) | 4 | JQ733706 | JQ733783 |
| *Dioscorea formosana* R.Knuth | Chen 42 (TCB) | 4 | JQ733691 | JQ733768 |
| *Dioscorea futschauensis* Uline ex R.Knuth | Hsu 32 (TCB) | 4 | JQ733671 | JQ733748 |
| *Dioscorea galeottiana* Kunth | Tellez 13090 (MEXU) | 1 | AY956499 | AY904796 |
| *Dioscorea gillettii* Milne-Redh. | Vollesen 61 (K) | 1 | AY956500 | AY904797 |
| *Dioscorea glabra* Roxb. | Wilkin et al. 874 (K) | 1 | AY956501 | AF307456 |
| *Dioscorea glomerulata* Hauman | Wood et al. 19466 (K) and Wood 8065 (K) | 5 | AY957586 | KM877894 |
| *Dioscorea gracilis* Hook. ex Poepp. | Villar & Viruel 18 (JACA) | 5 | KM877838 | KM877878 |
| *Dioscorea gracillima* Miq. | Tan 95288 (TNM) | 4 | JQ733709 | JQ733786 |
| *Dioscorea grandiflora* Mart. Ex Griseb. | Pereira 378 (K) | 5 | KM877850 | KM877895 |
| *Dioscorea hamiltonii* Hook.f. | Wilkin et al. 886 (K) | 1 | AY957587 | AF307465 |
| *Dioscorea hastifolia* Nees | Chase 2231 (K) | 5 | KM877868 | KM877932 |
| *Dioscorea hemsleyi* Prain & Burkill | Yang 18467 (TNM) | 4 | JQ733701 | JQ733778 |
| *Dioscorea hexagona* Baker | Wilkin et al. 960 (K) | 1 | AY957588 | AY904798 |
| *Dioscorea hispida* Dennst. | Wilkin et al. 855 (K) | 1 | AY957589 | AF307463 |
| *Dioscorea humifusa* Poepp. | Villar & Viruel 01 (JACA) | 5 | KM877839 | KM877879 |
| *Dioscorea humilis* subsp. *humilis* | Segarra-Moragues & Pérez-Collazos Eh01 (JACA) | 5 | KM877840 | KM877880 |
| *Dioscorea humilis* subsp. *polyanthes* (F.Phil.) Viruel, Segarra & Villar | Segarra-Moragues & Pérez-Collazos Ep01 (JACA) | 5 | KM877841 | KM877881 |
| *Dioscorea hyalinomarginata* Raz | Livshultz 1012 (BH) | 5 | KM877865 | KM877927 |
| *Dioscorea inopinata* Prain & Burkill | Thapayai et al. 513 (BKF) | 1 | AY957590 | AY904799 |
| *Dioscorea japonica* Thunb. | Hsu 94 (TCB) | 4 | JQ733665 | JQ733742 |
| *Dioscorea juxtalahuacensis* (O. Téllez & Dávila) Caddick & Wilkin | Calzada 21129 (MEXU) | 5 | KM877842 | KM877882 |
| *Dioscorea kamoonsensis* Kunth | CHC 7539 (TCB) | 4 | JQ733684 | JQ733761 |
| *Dioscorea karatana* Wilkin | Wilkin 950 (K) e Wilkin M947 (K) | 5 | AY957591 | KM877916 |
| *Dioscorea kituiensis* Wilkin & Muasya | Mwachala et al. 949A (EA) | 5 | KM877857 | KM877910 |
| *Dioscorea lanata* Bail | Miller 10423 (E) | 1 | AY957593 | AF307458 |
| *Dioscorea larecajensis* Uline ex R.Knuth | Wood 9960 (K) | 1 | AY957594 | newly produced |
| *Dioscorea lepcharum* Prain & Burkill | Lu 16156 (TCB) | 4 | JQ733675 | JQ733752 |
| *Dioscorea longirhiza* Caddick & Wilkin | Téllez 13081 (MEXU) | 5 | AY957595 | AF307473 |
| *Dioscorea maciba* Jum. & H. Perrier | Caddick et al. 318 (K) | 1 | AY957596 | AY904800 |
| *Dioscorea martini* Prain & Burkill | Yang 14136 (TNM) | 4 | JQ733710 | JQ733787 |
| *Dioscorea mayottensis* Wilkin | Hladik 8507 (P) | 5 | KM877861 | KM877917 |
| *Dioscorea mcvaughii* B.G. Schub. | Tellez 13080 (MEXU) | 1 | AY957597 | AF307460 |
| *Dioscorea melanophyma* Prain & Burkill | CHC 9203 (TCB) | 4 | JQ733688 | JQ733765 |
| *Dioscorea membranacea* Pierre ex Prain & Burkill | Wilkin et al. 878 (K) | 1 | AY957598 | AF307467 |
| *Dioscorea minima* C. B. Rob. & Seaton | Téllez 13086 (K) | 5 | KM877843 | KM877883 |
| *Dioscorea minutiflora* Engl. | Hladik 6235 (P) | 5 | KJ629250 | KM877933 |
| *Dioscorea modesta* Phill. | Villar & Viruel 06 (JACA) | 5 | KM877844 | KM877884 |
| *Dioscorea namorokensis* Wilkin | Wilkin et al. 1123 (K) e Wilkin 1126 (K) | 5 | AY957599 | KM877918 |
| *Dioscorea nipponica* Makino | Chase 6225 (K) | 1 | AY957600 | AF307455 |
| *Dioscorea nipponica* subsp. *rosthornii* (Diels) C. T. Ting | No data | 3 | HQ637585 | HQ637734 |
| *Dioscorea nitens* Prain & Burkill | Liang 2628 (TCB) | 4 | JQ733733 | JQ733810 |
| *Dioscorea nummularia* Lam. | Lu 20549 (TCB) | 4 | JQ733723 | JQ733800 |
| *Dioscorea orientalis* (J. Thiébaud) Caddick & Wilkin | Danin A (Silica specimen) | 5 | KM877858 | KM877911 |
| *Dioscorea oryzetorum* Prain & Burkill | Thapayai 464 (BKF) | 1 | AY972482 | AY904801 |
| *Dioscorea ovata* Vell. | Wood & Goyder 17012 (K) and Wood 9178 (K) | 5 | AY950686 | KM877900 |
| *Dioscorea pedicellata* Phil. | No data. | 5 | KM877845 | KM877886 |
| *Dioscorea pentaphylla* L. | Wilkin et al. 888 (K) | 1 | AY972483 | AF307470 |
| *Dioscorea petelotii* Prain & Burkill | Wilkin et al. 1023 (K) | 1 | AY973484 | AY904802 |
| *Dioscorea pohlii* Griseb. | Assis & Ladeira 67 (K) | 5 | KM877846 | KM877887 |
| *Dioscorea polystachya* Turcz. | Hsu 31 (TCB) | 4 | JQ733673 | JQ733750 |
| *Dioscorea prazeri* Prain & Burkill | Wilkin et al. 1075 (K) | 1 | AY973485 | AY904803 |
| *Dioscorea preussii* Pax | Davis 3019 (K) e Davi 2179 (K) | 5 | AY972486 | KM877920 |
| *Dioscorea pyrenaica* Bubani & Bordère ex Gren. | Segarra-Moragues & Catalán (Silica specimen) | 5 | KM877907 | KM877912 |
| *Dioscorea quinquelobata* Thunb. | Yonekura 6186 (TNM) | 4 | JQ733705 | JQ733782 |
| *Dioscorea rockii* Prain & Burkill | Wilkin et al. 1036 (K) | 1 | AY972488 | AY939882 |
| *Dioscorea sansibarensis* Pax | Wilkin et al. 973 (K) | 1 | AY972489 | AY939883 |
| *Dioscorea saxatilis* Poepp. | Villar & Viruel 11 (JACA) | 5 | KM877847 | KM877889 |
| *Dioscorea schimperiana* Hochst. ex Kunth | Wilkin et al. 762 (K) | 1 | AY972490 | AF307466 |
| *Dioscorea scorpioidea* C. Wright | Raz 339 (NY) | 5 | KM877866 | KM877928 |
| *Dioscorea scortechinii* Prain & Burkill | Lu 19238 (TCB) | 4 | JQ733714 | JQ733791 |
| *Dioscorea simulans* Prain & Burkill | Huang & Guo 0648583 (NAS) | 2 | EF614206 | EF614217 |
| *Dioscorea sinoparviflora* C. T. Ting, M. G. Gilbert & Turland | Huang & Guo 0648574 (NAS) | 2 | DQ974179 | DQ408171 |
| *Dioscorea soso* Jum. & H. Perrier | Wilkin et al. 1102 (K) and Wilkin et al. 1106 (K) | 1 | AY972491 | AY939884 |
| *Dioscorea spongiosa* J.Q. Xi, M. Mizuno & W.L.Zhao | Huang & Guo 0648581 (NAS) | 2 | DQ974191 | DQ974194 |
| *Dioscorea strydomiana* Wilkin | Lotter & Turpin 10627 (K) | 5 | KM877860 | KM877913 |
| *Dioscorea subcalva* Prain & Burkill | Liang 1045 (TCB) | 4 | JQ733682 | JQ733759 |
| *Dioscorea sylvatica* Eckl. | Chase 6184 (K) | 1 | AY972826 | AF307462 |
| *Dioscorea tentaculigera* Prain & Burkill | Thapayai 436 (BKF) | 1 | AY972828 | AY939886 |
| *Dioscorea tokoro* Makino ex Miyabe | Chen 5805 (TNM) | 4 | JQ733708 | JQ733785 |
| *Dioscorea tomentosa* J. König ex Spreng. | Chase 16311 (K) | 5 | KJ922833 | KM877925 |
| *Dioscorea trichantha* Baker | Wilkin et al. 1153 (K) | 1 | AY972829 | AY939887 |
| *Dioscorea variifolia* Betero | Villar & Viruel 03 (JACA) | 5 | KM877848 | KM877890 |
| *Dioscorea velutipes* Prain & Burkill | Liang 2609 (TCB) | 4 | JQ733732 | JQ733809 |
| *Dioscorea wallichii* Hook.f. | Wilkin et al. 1072 (K) | 1 | AY973830 | AY939888 |
| *Dioscorea yunnanensis* Prain & Burkill | Ma 7140 (TNM) | 4 | JQ733703 | JQ733780 |
| *Dioscorea zingiberensis* C.H. Wright | TCMK 244 (K) | 1 | AY973831 | AY939889 |
| *Dioscorea seriflora* Jum. & H. Perrier | Caddick et al. 302 | 1 | AY973827 | AY939885 |
| *Stemona tuberosa* Lour. | Hsu 401 (TCB) | 4 | JQ733738 | JQ733815 |
| *Stenomeris borneensis* Oliv. | Brun 19174 (K) and Lu 20601 (TCB) | 4 | AY973836 | JQ733811 |
| *Tacca artocarpifolia* Seem. | Caddick 305 (K) | 1 | newly<br>produced | AF307481 |
| *Tacca chantrieri* André | Chase 175 (NCU) | 1 | AY973837 | AJ286561 |
| *Tacca integrifolia* Ker Gawl. | Boyce 1074 (K) | 1 | AY973838 | AF307478 |
| *Tacca leontopetaloides* (L.) O. Kuntze | Wilkin et al. 817 (K) | 1 | AY973839 | AF307480 |
| *Tacca palmata* Blume | Boyce 1082 (K) | 1 | AY973840 | AF307479 |
| *Tacca palmatifida* Baker | Chase 1377 (K) | 1 | AY973841 | AY939890 |
| *Tacca plantaginea* (Hance) Drenth | Leiden 920520 (L) | 1 | AY973842 | AF307483 |
| *Thismia rodwayi* F. Muell. | Garnock-Jones 2218<br>(WELTU) | 1 | newly<br>produced | AY939892 |
| *Trichopus sempervivens* (H. Perrier)<br>Caddick & Wilkin | Caddick 304 (K) and<br>Wilkin 948 (K) | 1 | AF973844 | AF307476 |
| *Trichopus zeylanicus* Gaertn. | Caddick 346 (K) and<br>Chase 16354 (K) | 1 | AY973845 | AF307477 |

**Table S3.** Biogeographic models tested in this study using BioGeoBEARS package, and estimated parameters *d* (dispersion), *e* (extinction) and *j* (founder speciation event), log-likelihood and AIC values. Analysis performed on the clade New World I (a) and New World II (b).

**a) New World I**
| Model | LnL | d | e | j | ΔAICc | AICc weight |
| --- | --- | --- | --- | --- | --- | --- |
| DEC | -93.36 | 0.005779 | 1E-12 | 0 | 191.037 | 0.142 |
| DEC + J | -91.72 | 0.005095 | 1E-12 | 0.0150 | 190.094 | 0.228 |
| DIVALIKE | -97.93 | 0.007399 | 1E-12 | 0 | 200.176 | 0.0014 |
| DIVALIKE + J | -97.87 | 0.007373 | 1E-12 | 0.0001 | 200.072 | 0.0015 |
| BAYAREALIKE | -96.94 | 0.001960 | 4.30E-2 | 0 | 198.196 | 0.0039 |
| **BAYAREALIKE + J** | **-90.72** | **0.003768** | **1.00E-2** | **0.0333** | **188.093** | **0.6217** |

**b) New World II**
| Model | LnL | d | e | j | ΔAICc | AICc weight |
| --- | --- | --- | --- | --- | --- | --- |
| DEC | -46.53 | 0.007106 | 0.0025 | 0 | 97.875 | 0.0818 |
| DEC + J | -91.72 | 0.005095 | 1E-12 | 0.0150 | 191.160 | 4.531 |
| DIVALIKE | -45.94 | 0.008520 | 1.8E-9 | 0 | 96.698 | 0.147 |
| DIVALIKE + J | -49.11 | 0.008510 | 1E-12 | 1e-04 | 96.683 | 0.148 |
| BAYAREALIKE | -43.06 | 0.005526 | 0.0462 | 0.0462 | 103.022 | 0.0062 |
| **BAYAREALIKE + J** | **-43.06** | **0.003116** | **1E-07** | **1e-07** | **93.837** | **0.6160** |

**Table S4.**
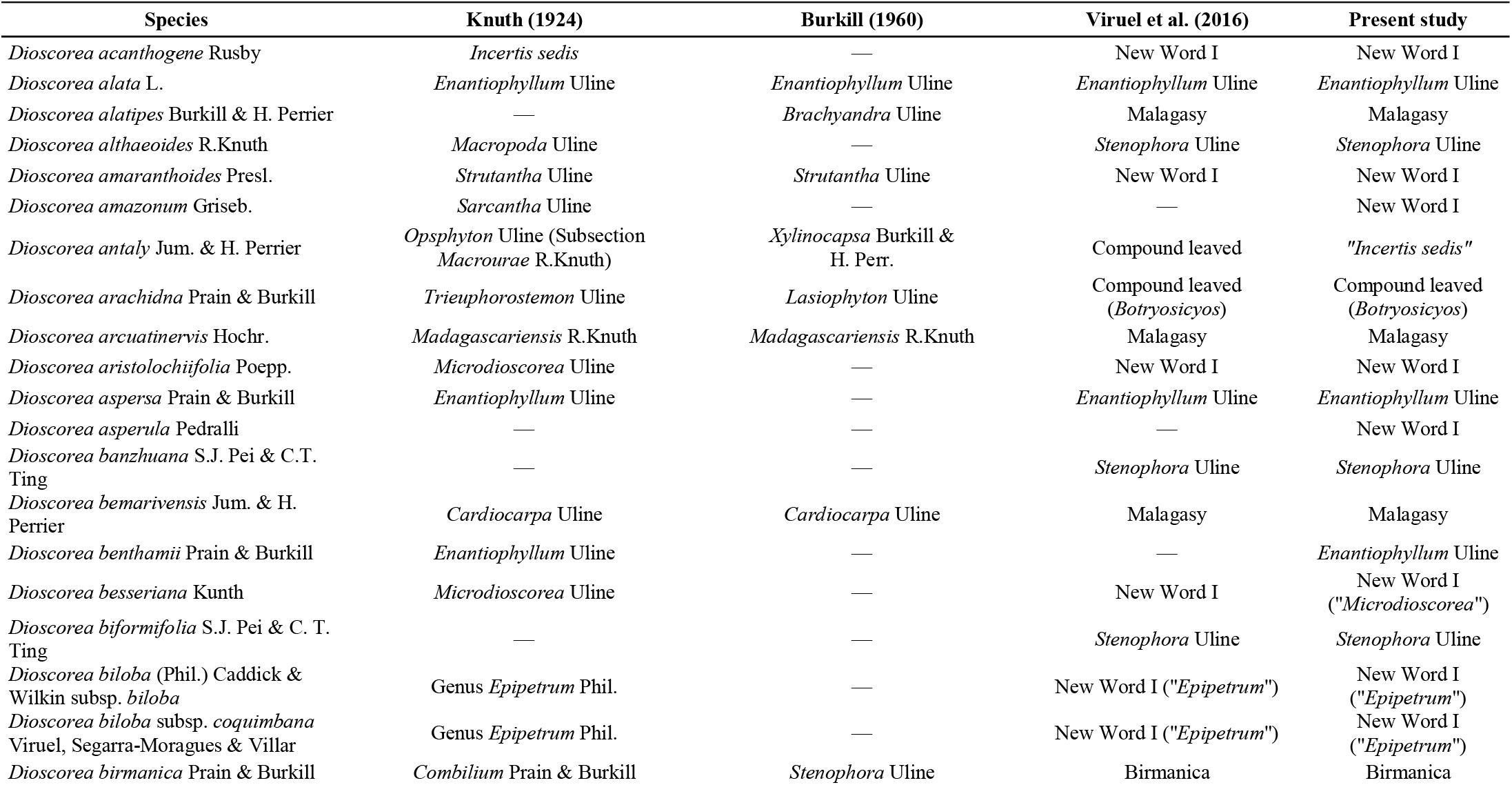
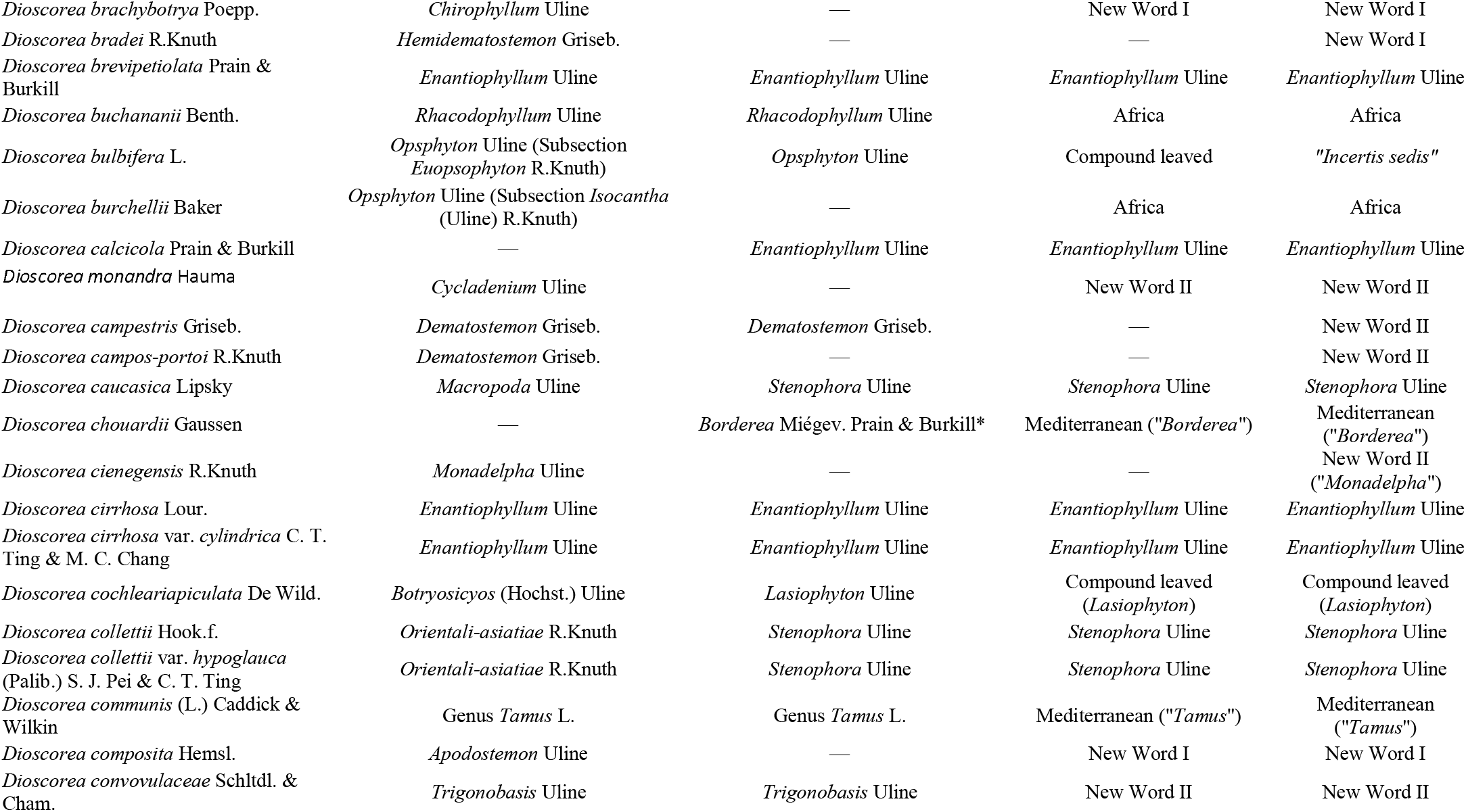
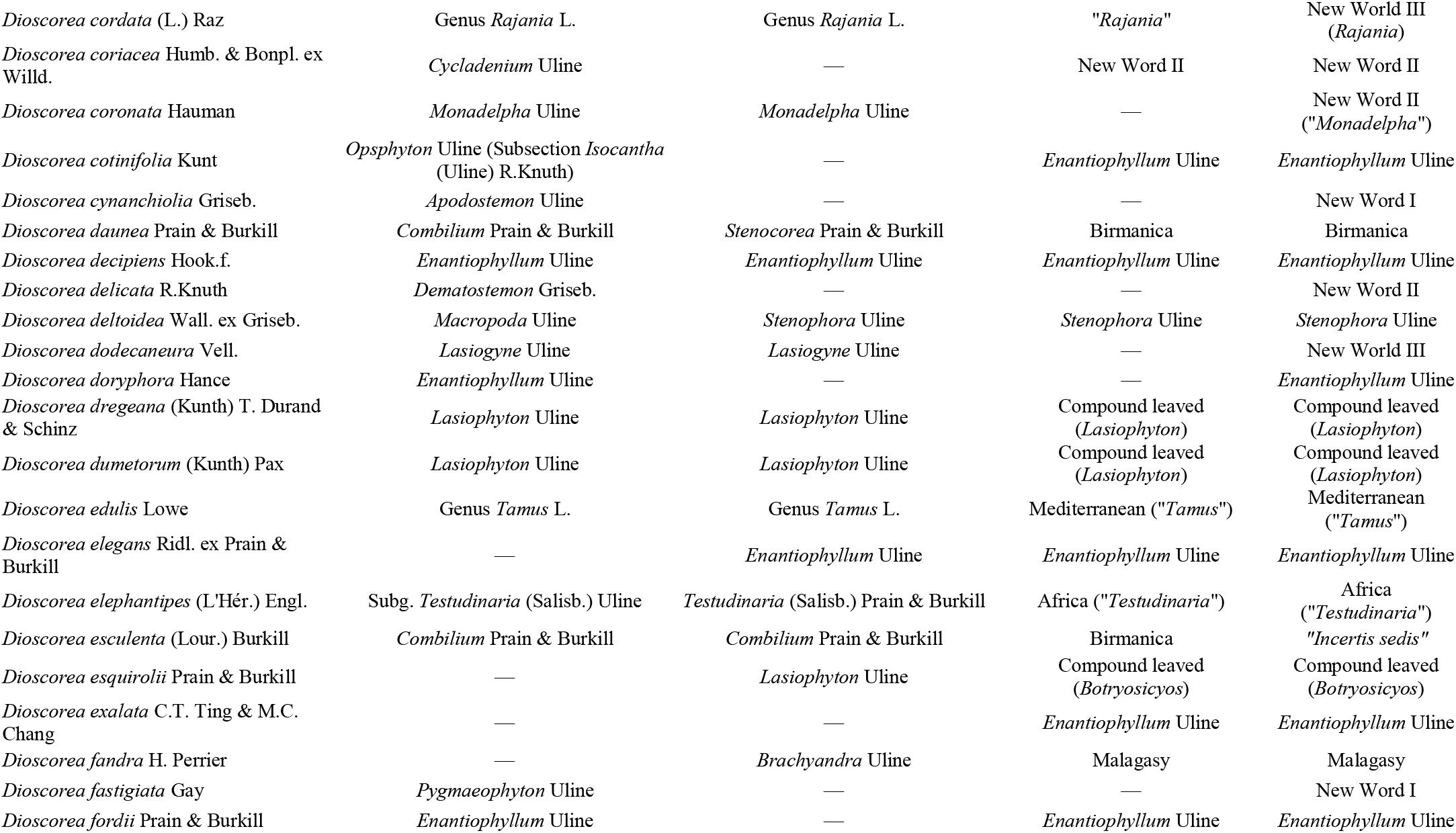
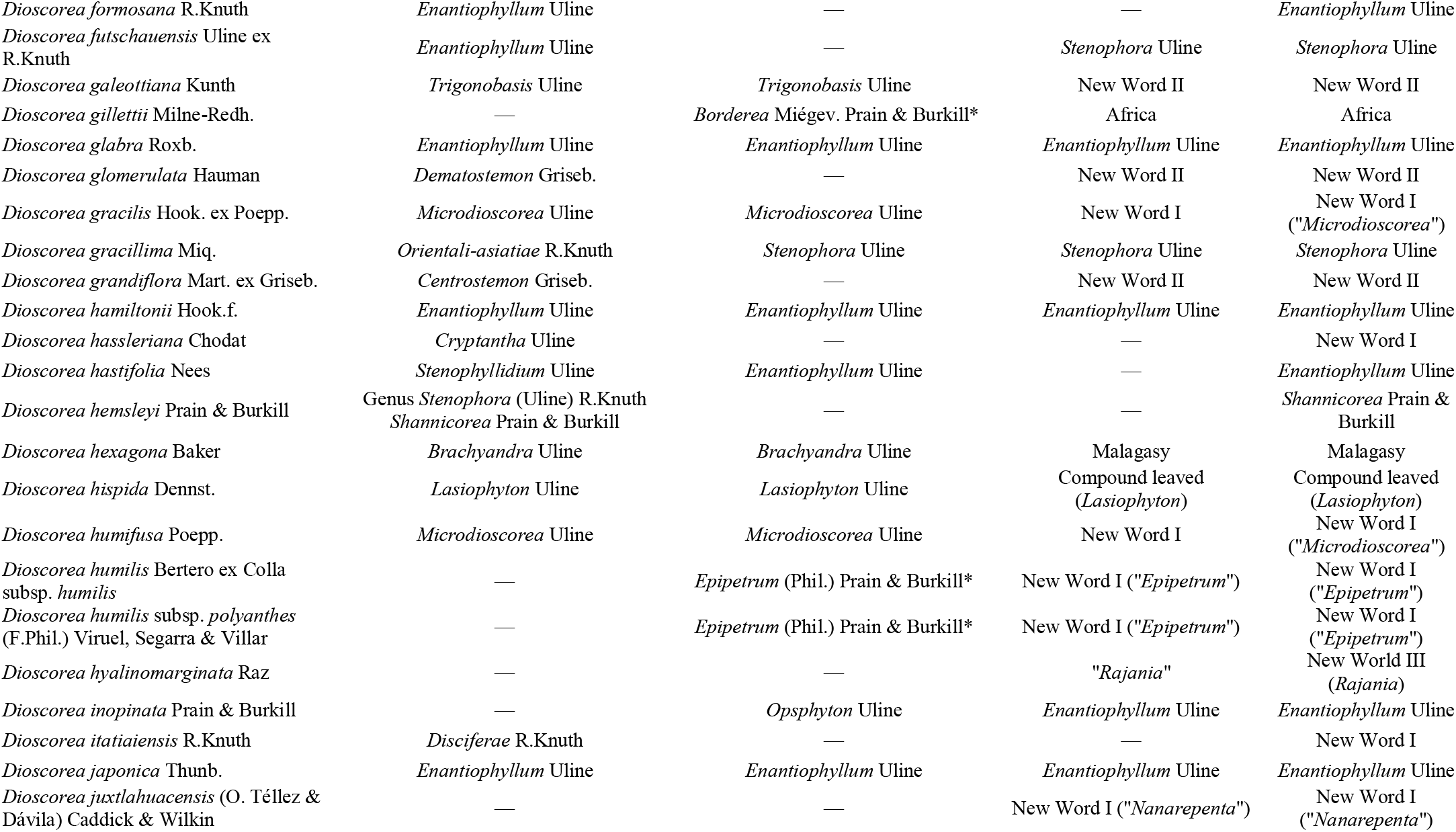
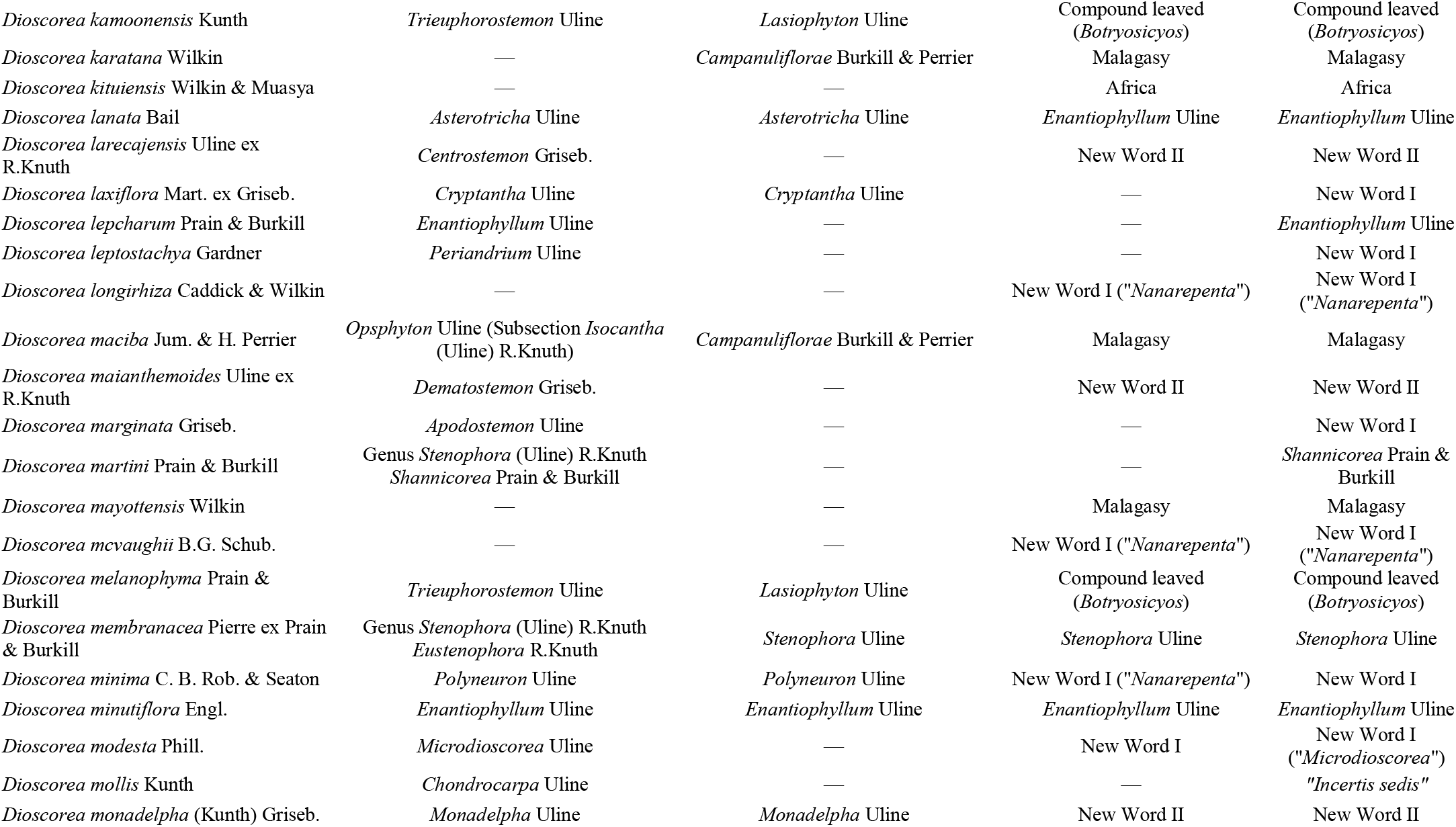
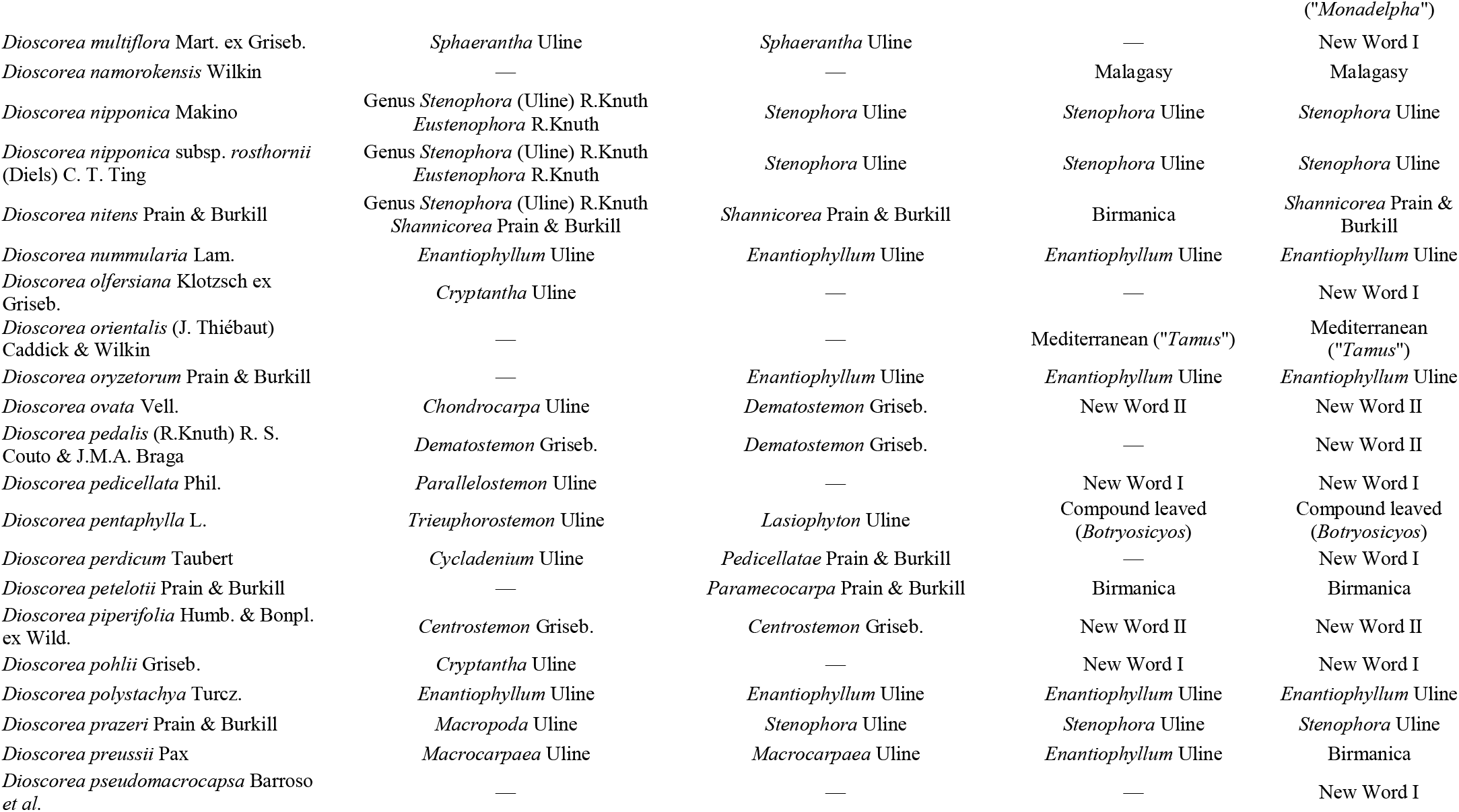
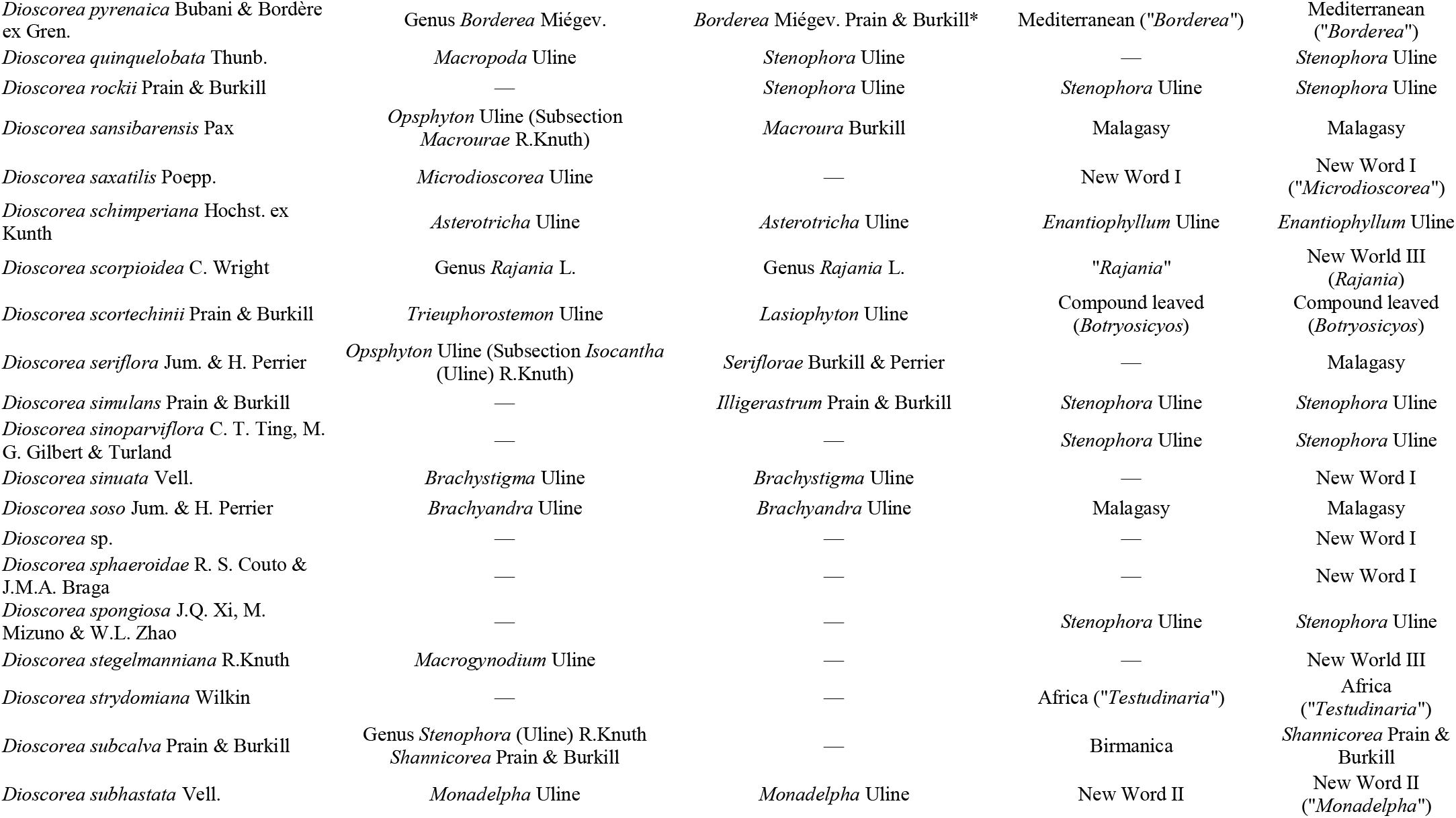
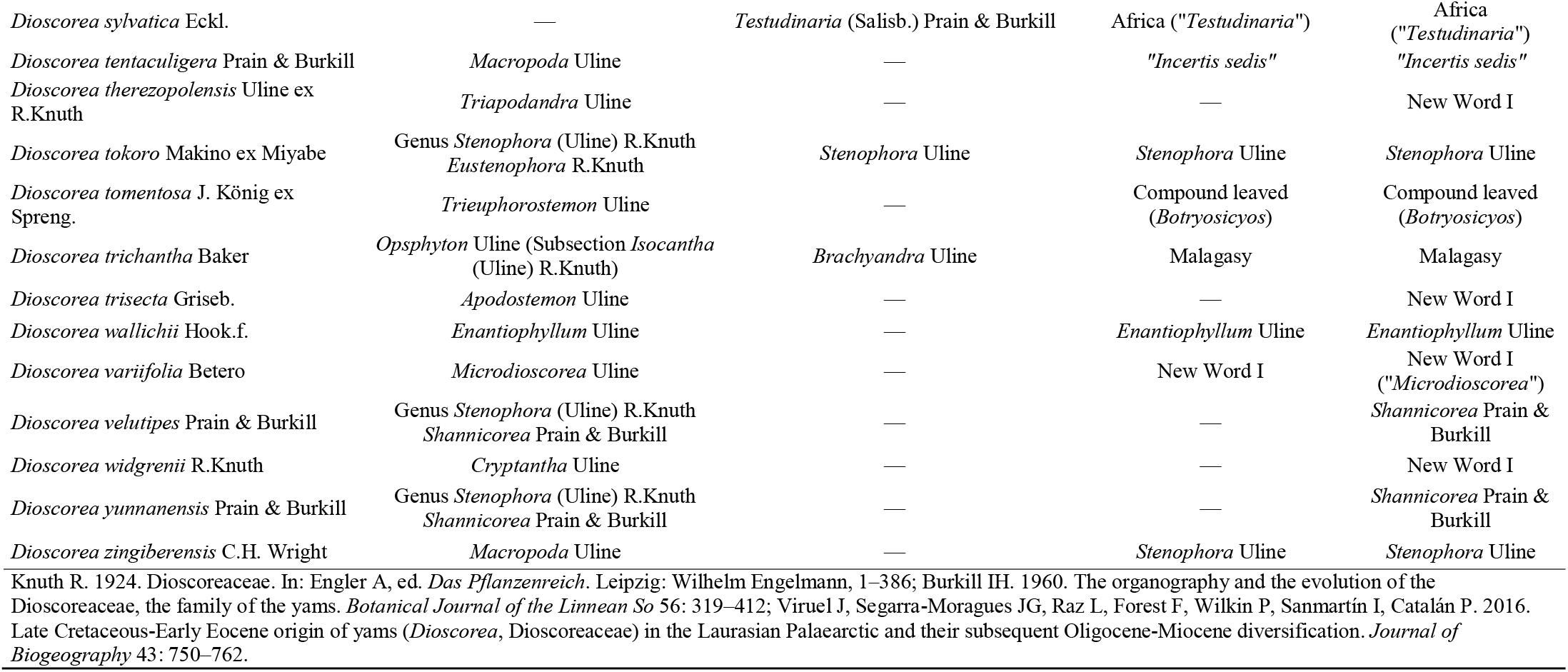
Species sampled and their position in two different infrageneric traditional classification systems for *Dioscorea* and in current phylogenetic molecular-based phylogenetic results. Complete references are given at the end.

**Table S5.** Species and their operational areas assigned in the biogeographic analysis.

| Species | Operational Areas |
| --- | --- |
| *Dioscorea amaranthoides* Presl. | D+E |
| *Dioscorea amazonum* Griseb. | D+E |
| *Dioscorea acanthogene* Rusby | D+E |
| *Dioscorea aristolochiifolia* Poepp. | C |
| *Dioscorea asperula* Pedralli | E |
| *Dioscorea besseriana* Kunth | C |
| *Dioscorea biloba* (Phil.) Caddick & Wilkin subsp. *biloba* | C |
| *Dioscorea biloba* subsp. *coquimbana* Viruel, Segarra-Moragues & Villar | C |
| *Dioscorea brachybotrya* Poepp. | C |
| *Dioscorea bradei* R.Knuth | F |
| *Dioscorea monandra* Hauman | F |
| *Dioscorea campestris* Griseb. | E+F |
| *Dioscorea campos-portoi* R.Knuth | E+F |
| *Dioscorea composita* Hemsl. | A |
| *Dioscorea convovulaceae* Schltld. & Cham. | A |
| *Dioscorea coriacea* Humb. & Bonpl. ex Willd. | B |
| *Dioscorea coronata* Hauman | F |
| *Dioscorea cynanchiolia* Griseb. | F |
| *Dioscorea delicata* R.Knuth | F |
| *Dioscorea fastigiata* Gay | C |
| *Dioscorea galeottiana* Kunth | A |
| *Dioscorea glomerulata* Hauman | C+F |
| *Dioscorea gracilis* Hook. ex Poepp. | C |
| *Dioscorea grandiflora* Mart. ex Griseb. | E+F |
| *Dioscorea hassleriana* Chodat | E+F |
| *Dioscorea humifusa* Poepp. | C |
| *Dioscorea humilis* Bertero ex Colla subsp. *humilis* | C |
| *Dioscorea humilis* subsp. *polyanthes* (F.Phil.) Viruel, Segarra & Villar | C |
| *Dioscorea* sp. | F |
| *Dioscorea itatiaiensis* R.Knuth | F |
| *Dioscorea juxtlahuacensis* (O. Téllez & Dávila) Caddick & Wilkin | A |
| *Dioscorea larecajensis* Uline ex R.Knuth | B |
| *Dioscorea laxiflora* Mart. ex Griseb. | D+E+F |
| *Dioscorea leptostachya* Gardner | E+F |
| *Dioscorea longirhiza* Caddick & Wilkin | A |
| *Dioscorea maianthemoides* Uline ex R.Knuth | E |
| *Dioscorea marginata* Griseb. | D+E+F |
| *Dioscorea mcvaughii* B.G. Schub. | A |
| *Dioscorea minima* C. B. Rob. & Seaton | A |
| *Dioscorea modesta* Phill. | C |
| *Dioscorea monadelphica* (Kunth) Griseb. | B+F |
| *Dioscorea multiflora* Mart. ex Griseb. | D+E+F |
| *Dioscorea olfersiana* Klotzsch ex Griseb. | F |
| *Dioscorea ovata* Vell. | F |
| *Dioscorea pedalis* (R.Knuth) R. S. Couto & J.M.A. Braga | F |
| *Dioscorea pedicellata* Phil. | C |
| *Dioscorea perdicum* Taubert | F |
| *Dioscorea piperifolia* Humb. & Bonpl. ex Wild. | D+E+F |
| *Dioscorea pohlii* Griseb. | D+E |
| *Dioscorea pseudomacrocapsa* Barroso *et al.* | F |
| *Dioscorea saxatilis* Poepp. | C |
| *Dioscorea cienegensis* R.Knuth | F |
| *Dioscorea sinuata* Vell. | F |
| *Dioscorea sphaeroidae* R. S. Couto & J.M.A. Braga | F |
| *Dioscorea subhastata* Vell. | F |
| *Dioscorea therezopolensis* Uline ex R.Knuth | F |
| *Dioscorea trisecta* Griseb. | E+F |
| *Dioscorea variifolia* Betero | C |
| *Dioscorea widgrenii* R.Knuth | F |

## REFERENCES

Antonelli A, Nylander JAA, Persson C, Sanmartín I. 2009. Tracing the impact of the Andean uplift on Neotropical plant evolution. Proceedings of the National Academy of Sciences of the United States of America 106: 9749–54.

Antonelli A, Sanmartín I. 2011. Mass extinction, gradual cooling, or rapid radiation? Reconstructing the spatiotemporal evolution of the ancient angiosperm genus Hedyosmum (Chloranthaceae) using empirical and simulated approaches. Systematic biology 60: 596–615.

Ayensu ES. 1972. Dioscoreales. In: Metcalfe CR, ed. Anatomy of the monocotyledons VI. Oxford: Clarendon Press,.

Ayensu ES., Coursey DG. 1972. Guinea Yams: the botany, ethnobotany, use and possible future of yams in West Africa. Economic Botany 26: 301–308.

Barroso GM, Sucre D, Guimarães EF, Carvalho LF, Valente MC, Silva JD, Silva JB, Rosenthal, F.R.T., Barbosa GM, Barth OM, Barbosa AF. 1974. Flora da Guanabara: família Dioscoreaceae. Sellowiana 25: 9–256.

Bolson M, De Camargo Smidt E, Brotto ML, Silva-Pereira V. 2015. ITS and trnH-psbA as Efficient DNA Barcodes to Identify Threatened Commercial Woody Angiosperms from Southern Brazilian Atlantic Rainforests. PLoS ONE 10 : e0143049.10.1371/journal.pone.0143049.

Burkill IH. 1960. The organography and the evolution of Dioscoreaceae, the family of the Yams. Journal of the Linnean Society of London, Botany 56: 319–412.

Caddick LR, Furness CA, Stobart KL, Rudall PJ. 1998. Microsporogenesis and pollen morphology in Dioscoreales and allied taxa. Grana 37: 321–336.

Caddick LR, Rudall PJ, Wilkin P, Hedderson TAJ, Chase MW. 2002a. Phylogenetics of Dioscoreales based on combined analyses of morphological and molecular data. Botanical Journal of the Linnean Society 138: 123–144.

Caddick LR, Wilkin P, Rudall PJ, Hedderson TAJ, Chase MW. 2002b. Yams reclassified: A recircumscription of Dioscoreaceae and Dioscoreales. Taxon 51: 103–114.

China Plant BOL. 2011. Comparative analysis of a large dataset indicates that internal transcribed spacer (ITS) should be incorporated into the core barcode for seed plants. PNAS 108: 19641–19646.

Choo T. 2009. Waging war against Dioscorea. Gardenwise: Newslett. Singapore Bot. Gard 32:17.

Christenhusz MJM, Chase MW. 2013. Biogeographical patterns of plants in the Neotropics - dispersal rather than plate tectonics is most explanatory. Botanical Journal of the Linnean Society 171: 277–286.

Costa LP. 2003. The historical bridge between the Amazon and the Atlantic forest of Brazil a study of molecular phylogeography with small mammals. Journal of Biogeography 30: 71–86.

Coursey DG. 1967. Yams: an account of the nature, origins, cultivation and utilisation of the useful members of the Dioscoreaceae. London: Longmans.

Couto RS, Tenorio V, Alzer F da C, Lopes RC, Vieira RC, Mendonça CBF, Gonçalves-Esteves V, Braga JMA. 2014. Taxonomic revision of the Dioscorea campestris species assemblage (Dioscoreaceae). Systematic Botany 39: 1056–1069.

Davis CC, Bell CD, Mathews S, Donoghue MJ. 2002. Laurasian migration explains Gondwanan disjunctions: evidence from Malpighiaceae. Proceedings of the National Academy of Sciences of the United States of America 99: 6833–7.

Davis CC, Webb CO, Wurdack KJ, Jaramillo CA, Donoghue MJ. 2005. Explosive radiation of Malpighiales supports a mid-Cretaceous origin of modern tropical rain forests. The American naturalist 165: 36–65.

Dorr LJ, Stergios B. 2003. A new species of Dioscorea (Dioscoreaceae) from the Andes of Venezuela. SIDA, contributions to Botany 20: 1007–1013.

Doyle JJ, Doyle JL. 1987. A rapid DNA isolation procedure from small quantities of fresh leaf tissue. Phytochemical Bulletin 19: 11–15.

Drummond AJ, Suchard MA, Xie D, Rambaut A. 2012. Bayesian phylogenetics with BEAUti and the BEAST 1.7. Molecular Biology and Evolution 29: 1969–73.

Dupin J, Matzke NJ, Särkinen T, Knapp S, Olmstead RG, Bohs L, Smith SD. 2017. Bayesian estimation of the global biogeographical history of the Solanaceae. Journal of Biogeography 44: 887–889.

e-Monocot team. 2017. The orders and families of monocots. Available at: http://e-monocot.org/ (accessed 30 August 2017).

Felsenstein J. 1985. Confidence-Limits on phylogenies - an approach using the bootstrap. Evolution 39: 783–791.

Freitas C, Meerow AW, Pintaud JC, Henderson A, Noblick L, Costa FRC, Barbosa CE, Barrington D. 2016. Phylogenetic analysis of Attalea (Arecaceae): insights into the historical biogeography of a recently diversified Neotropical plant group. Botanical Journal of the Linnean Society 182: 287–302.

Gao X, Zhu Y, Wu B, Zhao Y, Chen J, Hang Y. 2008. Phylogeny of Dioscorea sect. Stenophora based on chloroplast matK, rbcL and trnL-F sequences. Journal of Systematics and Evolution 46: 315–321.

Govaerts R, Wilkin P, Saunders RMK. 2007. World checklist of Dioscoreales: yams and their allies. Kew: Royal Botanical Gardens.

Gregor HJ. 1983. Erstnachweis der Gattung Tacca Forst 1776 (Taccaceae) im europäischen Alttertiär. Documenta Naturae 6: 27–31.

Hoorn C, Wesselingh FP, ter Steege H, Bermudez MA, Mora A, Sevink J, Sanmartin I, Sanchez-Meseguer A, Anderson CL, Figueiredo JP, Jaramillo C, Riff D, Negri FR, Hooghiemstra H, Lundberg J, Stadler T, Sarkinen T, Antonelli A. 2010. Amazonia through time: Andean uplift, climate change, landscape evolution, and biodiversity. Science 330: 927–931.

Hsu KM, Tsai LJ, Chen MY, Ku HM, Liu SC. 2013. Molecular phylogeny of Dioscorea (Dioscoreaceae) in East and Southeast Asia. Blumea 58: 21–27.

Hsu KM, Wang CM. 2012. Dioscorea sansibarensis Pax (Dioscoreaceae), a newly naturalized plant in Taiwan. Collection and Research 25: 25–29.

Knuth R. 1924. Dioscoreaceae. In: Engler A, ed. Das Pflanzenreich. Leipzig: Wilhelm Engelmann, 1–386.

Kress WJ, Erickson DL. 2007. A Two-Locus Global DNA Barcode for Land Plants: The Coding rbcL Gene Complements the Non-Coding trnH-psbA Spacer Region. PLoS ONE 2: 2007;2:e508.

Kunth KS. 1850. Enumeratio Plantarum Omnium Hucusque Cognitarum. : 908.

Landis MJ, Matzke NJ, Moore BR, Huelsenbeck JP. 2013. Bayesian analysis of biogeography when the number of areas is large. Systematic Biology 62: 789–804.

Levin RA, Wagner WL, Hoch PC, Nepokroeff M, Pires JC, Zimmer EA, Sytsma KJ. 2003. Family-level relationships of Onagraceae based on chloroplast rbcL and ndhF data. American Journal of Botany 90: 107–115.

Matuda E. 1961. Nuevas plantas de México. Anales del Instituto de Biología de la Universidad Nacional Autónoma de México 32: 143–147.

Matzke NJ. 2013. BioGeoBEARS: BioGeography with Bayesian (and Likelihood) Evolutionary Analysis in R Scripts.

Matzke NJ. 2014. Model selection in historical biogeography reveals that founder-event speciation is a crucial process in island clades. Systematic Biology 63: 951–970.

Maurin O, Muasya AM, Catalan P, Shongwe EZ, Viruel J, Wilkin P, Bank M Van Der. 2016. Diversification into novel habitats in the Africa clade of Dioscorea (Dioscoreaceae): erect habit and elephant’s foot tubers. BMC Evolutionary Biology 16: 238.

Mignouna HD, Abang MM, Geeta R. 2009. True Yams (Dioscorea): A Biological and Evolutionary Link between Eudicots and Grasses. Cold Spring Harbor Protocols 4(11): pdb.emo136.

Miller MA, Pfeiffer W, Schwartz T. 2010. Creating the CIPRES Science Gateway for inference of large phylogenetic trees. Gateway Computing Environments. New Orleans, USA, 1–8.

Morrone JJ. 2014. Biogeographical regionalisation of the Andean region. Zootaxa 3936 (2): 207–236.

O’Dea A, Lessios HA, Coates AG, Eytan RI, Restrepo-moreno SA, Cione AL, Collins LS, Queiroz A De, Farris DW, Norris RD, Stallard RF, Woodburne MO, Aguilera O, Aubry M pierre, Berggren WA, Budd AF, Cozzuol MA, Coppard SE, Duque-caro H, Finnegan S, Gasparini GM, Grossman EL, Johnson KG, Keigwin LD, Knowlton N, Leigh EG, Leonard-pingel JS, Marko PB, Pyenson ND, Rachello-dolmen PG, Soibelzon E, Soibelzon L, Todd JA, Vermeij GJ, Jackson JBC. 2016. Formation of the Isthmus of Panama. Science Advances 2: 1–12.

Olmstead RG. 2013. Phylogeny and biogeography in Solanaceae, Verbenaceae and Bignoniaceae: A comparison of continental and intercontinental diversification patterns. Botanical Journal of the Linnean Society 171: 80–102.

Pan AD, Jacobs BF, Currano ED. 2014. Dioscoreaceae fossils from the late Oligocene and early Miocene of Ethiopia. Botanical Journal of the Linnean Society 175: 17–28.

Philippi RA. 1864. Plantarum novarum Chilensiam Centuriae, inclusis quibusdam Mendosis et Patagoniois. Linnaea 33: 253.

Potonié H. 1921. Lehrbuch der Paleobotanik. Berlin: Gebrüder Bornträger.

Prain D, Burkill IH. 1914. A Synopsis of the Dioscorea of the Old World, Africa excluded, with description of new species and of varieties. J. Proc. Asiat. Soc. Bengal 10: 5–41.

R Core Team. 2016. R: a language and environment for statistical computing. R Foundation for Statistical Computing, Vienna, Austria. Available at: www.R-project.org/. (accessed 30 August 2015).

Rambaut A. 2009. FigTree: Tree figure drawing tool. Available at: http://tree.bio.ed.ac.uk/software/figtre?e/. (accessed 30 August 2017).

Rambaut A, Suchard MA, Drummond AJ. 2014. Tracer v1.6, 2003-2013: MCMC trace analysis tool. Available at: http://tree.bio.ed.ac.uk/software/tracer/. (accessed 30 August 2017).

Raz L. 2002. Dioscoreaceae in Flora of North America (efloras). Available at: http://www.efloras.org/florataxon.aspx?flora_id=1&taxon_id=10280. (accessed 30 August 2017).

Raz L. 2016. Untangling the West Indian Dioscoreaceae: new combinations, lectotypification and synonymy. Phytotaxa 258: 26–48.

Raz L. 2017. A review of the fossil record of Dioscoreaceae. Botanical Journal of the Linnean So 183: 495–508.

Raz L., Pérez-Camacho J. 2016. A new species of Dioscorea (Dioscoreaceae) from Central Cuba. Brittonia 69: 109–113.

Ree RH, Smith SA. 2008. Maximum likelihood inference of geographic range evolution by dispersal, local extinction, and cladogenesis. Systematic Biology 57: 4–14.

Renner S. 2004. Plant dispersal across the tropical atlantic by wind and sea currents. International Journal of Plant Sciences 165: S23–S33.

Ronquist F. 1997. Dispersal-vicariance analysis: a new approach to the quantification of historical biogeography. Systematic Biology 46: 195.

Ronquist F, Teslenko M, van der Mark P, Ayres DL, Darling A, Höhna S, Larget B, Liu L, Suchard MA, Huelsenbeck JP. 2012. MrBayes 3.2: efficient Bayesian phylogenetic inference and model choice across a large model space. Systematic Biology 61: 539–42.

Rouhan G, Labiak PH, Randrianjohany E, Rakotondrainibe F. 2012. Not so Neotropical after all: the grammitid fern genus Leucotrichum (Polypodiaceae) is also paleotropical, as revealed by a new species from Madagascar. Systematic Botany 37: 331–338.

Schols P, Furness CA, Wilkin P, Huysmans S, Smets E. 2001. Morphology of pollen and orbicules in some Dioscorea species and its systematic implications. Botanical Journal of the Linnean Society 136: 295–311.

Schols P, Furness CA, Wilkin P, Smets E, Cielen V, Huysmans S. 2003. Pollen morphology of Dioscorea (Dioscoreaceae) and its relation to systematics. Botanical Journal of the Linnean Society 143: 375–390.

Silvestro D, Michalak I. 2012. raxmlGUI: a graphical front-end for RAxML. Organisms Diversity & Evolution 12: 335–337.

Simon MF, Grether R, de Queiroz LP, Skema C, Pennington RT, Hughes CE. 2009. Recent assembly of the Cerrado, a neotropical plant diversity hotspot, by in situ evolution of adaptations to fire. Proceedings of the National Academy of Sciences 106: 20359–20364.

Staden R, Judge DP, Bonfield JK. 2003. Analysing sequences using the Staden Package and EMBOSS. In: Krawetz SA,, In: Womble DD, eds. Introduction to Bioinformatics: A theoretical and practical approach. Totawa: Humana Press,.

Swofford DL. 2002. PAUP*. Phylogenetic Analysis Using Parsimony (*and Other Methods). Sinauer Associates, Sunderland, Massachusetts.

Tamura KS, Stecher G, Peterson D, Filipski A. 2013. MEGA6: Molecular evolutionary genetics analysis version 6.0. Molecular Biology and Evolution 30: 2725–2729.

Téllez-Valdés O, Dávila-Aranda P. 1998. Nanarepenta juxtlahuacensis (Dioscoreaceae), una nueva especie de Oaxaca, México. Novon 8: 210–214.

Tenorio V, Couto RS, Albuquerque ESB, Braga JMA, Vieira RC. 2017. Stem anatomy of neotropic Dioscorea L. (Dioscoreaceae) and its importance to the systematics of the genus. Plant Systematic and Evolution: 303(6): 775–786.

The Plant List. 2013. The Plant List: a working list of all plant species. Available at: http://www.theplantlist.org/ (accessed 30 August 2017)

Thiers B. Index Herbariorum: A global directory of public herbaria and associated staff. New York Botanical Garden’s Virtual Herbarium.

Thompson JD, Gibson TJ, Plewniak F, Jeanmougin J, Higgins DG. 1997. The ClustalX windows interface: flexible strategies for multiple sequence alignment aided by quality analysis tools. Nucleis Acids Research 24: 4876–4882.

Uline EH. 1897. Dioscoreaceae. In: Engler, A. & Prantl K, ed. Die Naturlichen Pflanzenfamilien. Leipzig: Engelmann, 80–87.

Viruel J, Segarra-Moragues, J.G., Pérez-Collazos E, Villar L, Catalán P. 2010. Systematic revision of the Epipetrum group of Dioscorea (Dioscoreaceae) endemic to Chile. Systematic Botany 35: 40–63.

Viruel J, Segarra-Moragues JG, Raz L, Forest F, Wilkin P, Sanmartín I, Catalán P. 2016. Late Cretaceous-Early Eocene origin of yams (Dioscorea, Dioscoreaceae) in the Laurasian Palaearctic and their subsequent Oligocene-Miocene diversification. Journal of Biogeography 43: 750–762.

Werneck FP, Nogueira C, Colli GR, Sites JW, Costa GC. 2012. Climatic stability in the Brazilian Cerrado: Implications for biogeographical connections of South American savannas, species richness and conservation in a biodiversity hotspot. Journal of Biogeography 39: 1695–1706.

Wilkin P, Muasya AM, Banks H, Furness CA, Vollesen K, Weber O, Demissew S. 2009. A new species of yam from Kenya, Dioscorea kituiensis: pollen morphology, conservation status, and speciation. Systematic Botany 34: 652–659.

Wilkin P, Schols P, Chase MW, Chayamarit K, Furness CA, Huysmans, S., Rakotonasolo F, Smets E, Thapyai C. 2005. A plastid gene phylogeny of the yam genus, Dioscorea: roots, fruits and Madagascar. Systematic Botany 30: 736–749.

Zachos JC, Dickens GR, Zeebe RE. 2008. An early Cenozoic perspective on greenhouse warming and carbon-cycle dynamics. Nature 451: 279–283.

